# Tau phosphorylation impedes functionality of protective tau envelopes

**DOI:** 10.1101/2024.03.25.586522

**Authors:** Valerie Siahaan, Romana Weissova, Eva Lanska, Adela Karhanova, Vojtech Dostal, Veronique Henriot, Carsten Janke, Lenka Libusova, Marcus Braun, Martin Balastik, Zdenek Lansky

## Abstract

Tau, an axonal microtubule-associated protein, is a critical regulator of microtubule function and stability. Tau interaction with microtubules is regulated by tau phosphorylation. Tau hyperphosphorylation is implicated in microtubule destabilization related to neurodegenerative disorders. How tau phosphorylation leads to microtubule destabilization is however unknown. Recently, it was shown that tau molecules on microtubules cooperatively assemble into cohesive layers termed envelopes. Tau envelopes protect microtubules against degradation by microtubule-severing enzymes, suggesting a functional link between envelopes and microtubule stability. Here we show that tau phosphorylation has deleterious effects on the microtubule-protective function of tau envelopes. Using reconstitution and live-cell experiments, we found that tau phosphorylation destabilizes tau envelopes and decreases their integrity, leading to reduced microtubule protection against microtubule-severing enzymes. Our data suggest that a perturbation of microtubule homeostasis linked to tau hyperphosphorylation in neurodegeneration, could be explained by the disassembly and impaired functionality of the tau envelopes.

## Introduction

Microtubules are rigid filaments, which are essential for neuronal function and homeostasis e.g. by providing tracks for intracellular cargo transport. Microtubules are dynamic polymers, frequently switching between phases of assembly and disassembly^1^. The lifetime of a microtubule is regulated by a multitude of microtubule-associated proteins, which either affect microtubule assembly and disassembly^2,3^, or can sever microtubules into fragments^4^. Deregulation of various microtubule-associated proteins, e.g. tau, has been shown to trigger changes in microtubule dynamics, induce loss of microtubule mass from axons and dendrites, and is associated with multiple neurodegenerative disorders^2,5^.

Tau is an intrinsically disordered microtubule-associated protein, which in healthy neurons localizes predominantly to axonal microtubules. Tau regulates the functioning of other microtubule-associated proteins and protects microtubules against microtubule-severing enzymes, such as katanin^6^. During neurodegeneration, tau is found aggregated in neurofibrillary tangles, which is one of the hallmarks of neurodegenerative disorders collectively termed tauopathies, such as Alzheimer’s disease^7,8^. It was proposed that tau aggregation causes a depletion of functional tau^9,10^, thereby leaving the axonal microtubules unprotected against microtubule-severing enzymes, such as katanin, which could lead to pathological microtubule destabilization. Moreover, aggregated tau has an increased phosphorylation state as compared to physiological tau, and has been shown to have reduced interaction with microtubules^11,12^. These findings suggest a relation between hyperphosphorylation of tau and microtubule instability related to neurodegeneration, nevertheless, the underlying molecular mechanism remains unclear.

It has recently been shown that tau molecules associate with microtubules in two distinct modes – either (i) diffusing individually along the microtubule lattice, rapidly binding and unbinding, or (ii) binding cooperatively, with much longer interaction times, constituting cohesive envelopes, previously referred to as ‘condensates’ or ‘islands’^13–15^. These envelopes enclose the microtubules and act as selectively permeable barriers for other microtubule-associated proteins^13,14^. Tau envelopes can differentially modulate the action of microtubule-related molecular motors, e.g. decreasing the kinesin-1 walking distance^14,16^, while permitting dynein-mediated transport^13^. While microtubules covered by individually diffusing tau molecules are prone to disintegration by microtubule severing enzymes, such as katanin^14^, tau envelopes efficiently protect the microtubule surface from the action of microtubule severing enzymes^13,14^. These observations suggest that the protective function of tau is mediated by the cohesion of tau envelopes. We thus hypothesized that pathological effects of tau phosphorylation can be explained by the impact of tau phosphorylation on the formation and function of protective tau envelopes.

Here, we demonstrate that the formation and maintenance of tau envelopes is indeed critically regulated by phosphorylation. We found that phosphorylation of tau decreases the propensity of tau to form envelopes and that envelopes formed by phosphorylated tau have altered functionality with decreased protection against microtubule severing enzymes. Our findings suggest that the cohesive binding mode of tau may provide a causal connection between tau phosphorylation and impaired tau functionality: the reduction of tau envelopes and their impaired functionality, caused by tau phosphorylation, results in decreased shielding of microtubules from severing enzymes, and consequently in a decrease of microtubule stability.

## Results

### Tau phosphorylation induces envelope disassembly

To investigate whether phosphorylation of tau affects the formation of tau envelopes, we expressed GFP-labelled human 2N4R tau (full length tau protein, 441 amino acids), in insect cells and used Alkaline phosphatase to dephosphorylate the tau in vitro (Methods). This approach yielded two tau samples: phosphorylated tau (native, insect cell expressed tau, denoted as ‘phospho-tau’), and dephosphorylated tau (phosphatase treated, insect cell expressed tau, denoted as ‘dephospho-tau’) (Fig. 1a). To confirm the efficiency of the phosphatase-treatment, we determined the degree of phosphorylation of these samples at all potential phosphorylation sites using mass spectrometry (Methods, Fig. 1b, for individual sites see Supplementary Fig. 1a,b). We then added the phospho-tau or dephospho-tau samples at 1.5 nM to surface-immobilized taxol-stabilized microtubules and visualized the interaction using TIRF microscopy. While dephospho-tau readily formed micrometer-sized envelopes at this concentration, by contrast, phospho-tau was present on microtubules only diffusively and did not form envelopes (Fig. 1c, Supplementary Movie 1,2). Repeating this experiment at two higher concentrations of tau, we found that tau envelopes were formed by both tau samples (Supplementary Fig. 1c), nevertheless, at all concentrations tested, dephospho-tau covered a higher percentage of the microtubules compared to phospho-tau (Fig. 1d), demonstrating higher propensity of dephosphorylated tau to form envelopes. To confirm our findings, we repeated these experiments with tau expressed in bacterial cells which possesses a low phosphorylation state (denoted by Bact-tau) and used a kinase to increase its phosphorylation state (Supplementary Fig. 1d, Methods). Multitude of kinases have been shown to phosphorylate tau, including proline-directed kinases (e.g. Cdk5, GSK-3β or MAP kinases). Cdk5-mediated phosphorylation of tau has been shown in healthy conditions to control multiple processes in neural development (e.g. axonal growth and guidance), while hyperactivation of Cdk5 (e.g. in Alzheimer’s disease) has been shown to result in heightened tau phosphorylation, promoting tau mislocalization, aggregation and formation of neurofibrillary tangles^9,17,18^. Therefore, we phosphorylated the Bact-tau sample using Cdk5 kinase with its activator p35 (denoted by Bact-Cdk5-tau, Supplementary Fig. 1d, Methods) yielding a sample with higher phosphorylation degree as confirmed by mass spectrometry (Supplementary Fig. 1e, for individual sites see Supplementary Fig. 1f,g). We then added these samples separately to surface-immobilized taxol-stabilized microtubules and studied the envelope coverage after 3 minutes of incubation. In accordance with our previous results, we found that the Bact-tau formed envelopes at much lower concentrations compared to Bact-Cdk5-tau (Supplementary Fig. 1h). We next asked if tau phosphorylation can destabilize preexisting tau envelopes formed by dephosphorylated tau. To test this, we formed envelopes using 15 nM Bact-tau and after 10 minutes we added active Cdk5 kinase to the channel while keeping tau in solution. After the addition of active kinase we observed that the tau envelopes started to disassemble from their boundaries (Fig. 1e, Supplementary movie 3, Methods) with occasional fission events within the boundaries of the envelope during disassembly (0.01 ± 0.08 fissions mm^-1^s^-1^). In a control experiment, we added deactivated Cdk5 (Methods) to the envelopes while keeping tau in solution, in which case no disassembly was observed and, on the contrary, a significant increase of the envelope coverage was detected (Fig. 1e,f, Supplementary movie 4). Combined, these experiments show that phosphorylation of tau decreases the propensity of tau to form envelopes and destabilizes preexisting envelopes.

**Fig. 1.**
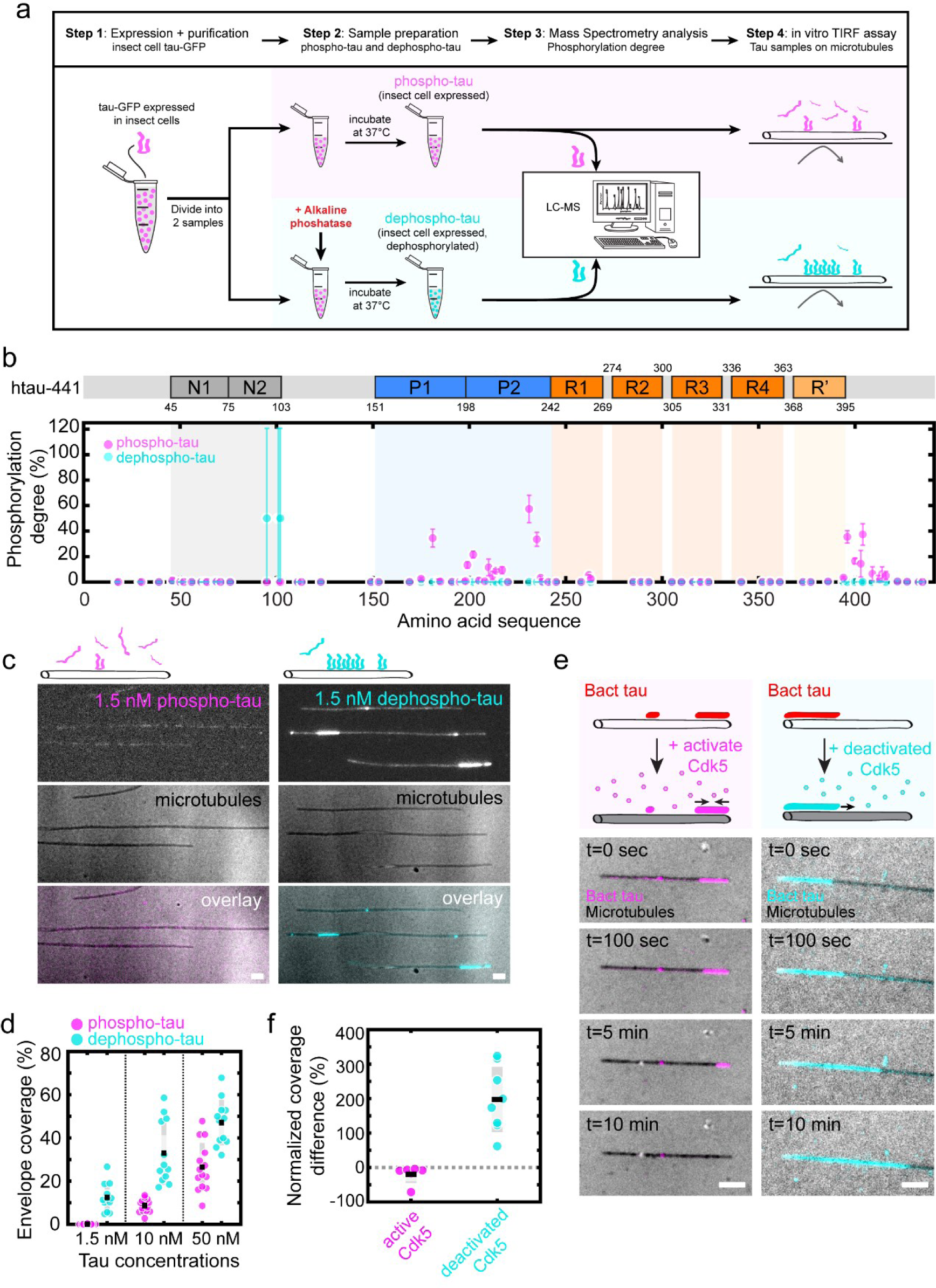
Tau phosphorylation induces envelope disassembly. **a**. Schematics of the sample preparation. **b**. Mass-spectrometry-determined degree of phosphorylation of phospho-tau (insect cell expressed tau, magenta) and dephospho-tau (phosphatase-treated insect cell expressed tau, cyan). Phosphorylation degree is presented as the mean ± s.d. (Methods) and displayed at the location of the phosphorylation site along the amino acid sequence of tau (schematic of the sequence is shown above the plot). The domains on the tau sequence are color-coded: N-terminal domains (N1, N2, grey), proline-rich domains (P1, P2, blue), microtubule-binding repeats (R1-R4, orange), and the domain pseudo-repeat (R‘, light orange). **c**. Multichannel fluorescence micrographs of 1.5 nM phospho-tau (magenta, left), and 1.5 nM dephospho-tau (cyan, right) on taxol-stabilized microtubules (black, middle panels) after 3 min incubation. Scale bars: 2 μm. **d**. Percentage of taxol-stabilized microtubules covered with tau envelopes after 3 min of tau incubation on surface-immobilized microtubules. Envelope coverage for phospho-tau (magenta) at 1.5 nM was 0.1 ± 0.1%, at 10 nM was 8.6 ± 3.1%, and at 50 nM was 26.5 ± 11.2% (mean ± s.d., n=14, 14, 14 fields of view in 11 independent experiments). Envelope coverage for dephospho-tau (cyan) at 1.5 nM was 12.5 ± 6.4%, at 10 nM was 32.9 ± 14.7%, and at 50 nM was 47.1 ± 10.5%, n=12, 12, 12 independent experiments). **e**. Multichannel fluorescence micrographs of 15 nM Bact-tau (red in schematics) after treatment with active Cdk5 (left, Bact-tau in magenta) or with deactivated Cdk5 (right, Bact-tau in cyan). Microtubules (black) imaged using IRM. Scale bars: 2μm. **f.** Normalized difference between the coverage before and after treatment with active Cdk5 or deactivated Cdk5. Normalized coverage difference for active Cdk5: −19.4 ± 25.9% (mean ± s.d., n=6 independent experiments); deactivated Cdk5: 197.6 ± 94.3% (n=8 independent experiments). Two-sided t-test: p=1.52*10^-4^.

### Phosphorylation reduces envelope integrity

We next asked why tau phosphorylation leads to a decrease in envelope formation. We hypothesized that phosphorylated tau might not be able to participate in the formation of tau envelopes. Our phosphorylated tau samples are heterogeneous, meaning that they consist of tau molecules with different patterns and degrees of phosphorylation, which could be differently competent in forming envelopes. To test which tau molecules, out of a given sample, can participate in envelope formation, we added phospho-tau at high concentration (1 μM) to microtubules in solution, ensuring that microtubules were fully covered by tau envelopes. By spinning down the microtubules, we then separated the sample into (i) tau, which participated in envelope formation (tau in envelopes – characterized by limited translocation along the microtubule lattice and with very low turnover^13,14^ - found in the pellet along with the microtubules) and (ii) tau, which did not participate in envelope formation (tau in solution – unbound tau or tau characterized by rapid diffusion along the microtubule lattice and high turnover^13,14^ - found in the supernatant) (Fig. 2a, Methods). When we then analyzed the phosphorylation degree of these two samples by mass spectrometry, we did not find any striking differences in the phosphorylation patterns between the two samples (Fig. 2b, for individual sites see Supplementary Fig. 2a,b), demonstrating that phosphorylated tau is competent in envelope formation. We did, nevertheless, find that the phosphorylation degree of tau in the supernatant (not participating in envelope formation) was slightly elevated compared to tau in the envelopes. While we observed almost no phosphorylation in the N-terminal region as well as the microtubule-binding repeats for both tau samples, we found on average about a 5% increase in phosphorylation degree within the proline-rich region and C-terminal regions in tau found in solution. This data demonstrates that although phosphorylation of tau decreases its propensity to form envelopes, phosphorylated tau can (particularly at higher concentration) be incorporated into the tau envelopes.

**Fig. 2.**
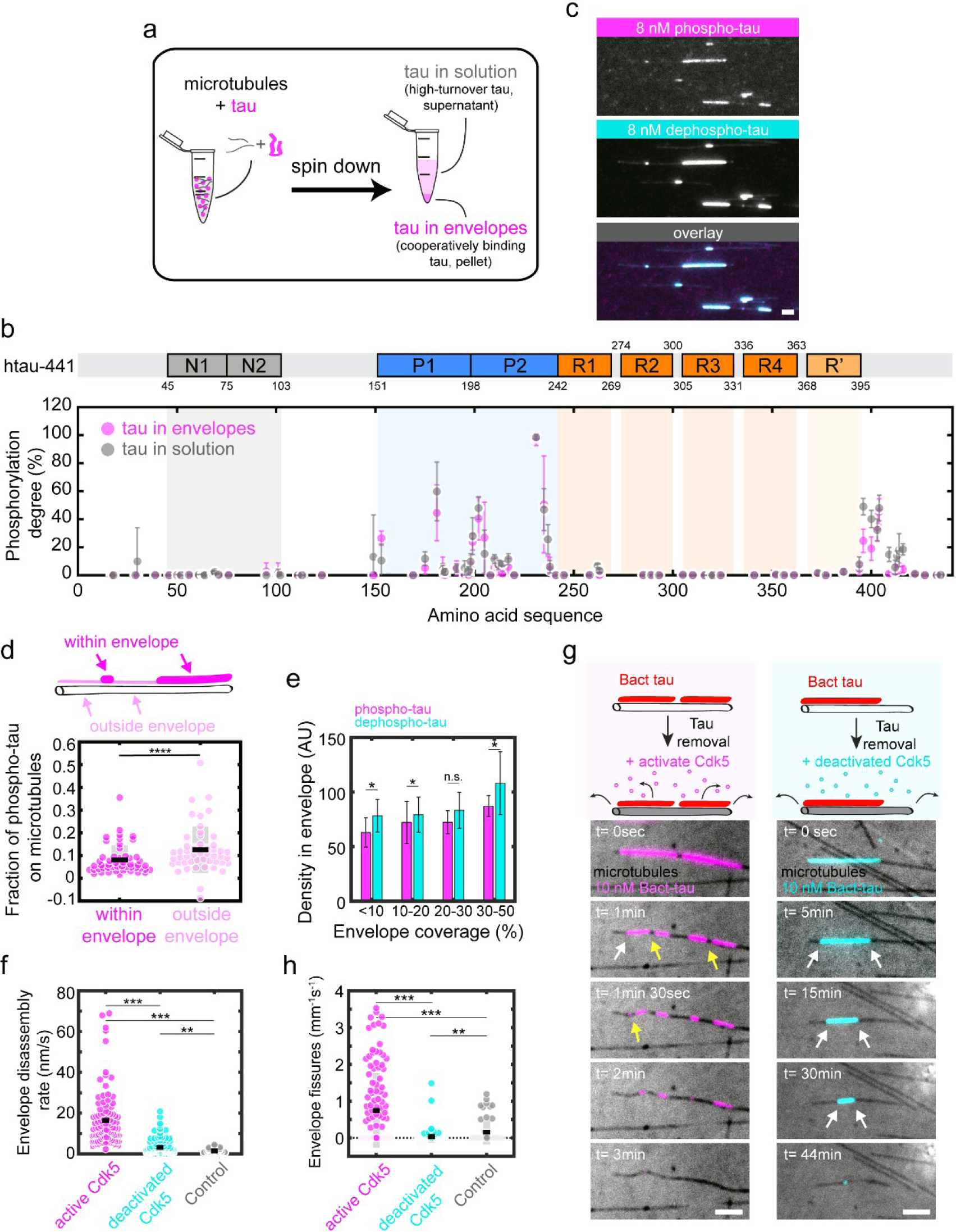
Phosphorylation reduces envelope integrity. **a**. Schematics of the sample preparation (Methods). **b**. Mass-spectrometry-determined degree of phosphorylation of cooperatively bound tau found in the pellet (slow-turnover tau in envelopes, magenta) and unbound tau found in the supernatant (high-turnover tau in solution, grey). The phosphorylation degree was presented as the mean ± s.d. for each sample (Methods) and displayed at the location of the phosphorylation site along the amino acid sequence of tau (schematic of the sequence is shown above the plot). The domains on the tau sequence are color-coded: N-terminal domains (N1, N2, grey), proline-rich domains (P1, P2, blue), microtubule-binding repeats (R1-R4, orange), and the domain pseudo-repeat (R‘, light orange). **c**. Multichannel fluorescence micrographs of 8 nM phospho-tau (magenta, top panel) and 8 nM dephospho-tau (cyan, middle panel) incubated simultaneously (overlay, bottom panel) and imaged after 3 min incubation. Scale bar: 2 μm. **d.** Fraction of the density of phospho-tau compared to the total density of all tau within tau envelope region (magenta), or outside of tau envelope region (light pink). Location of the envelope and non-envelope region are schematically drawn in the cartoon above the plot. Fraction of phospho-tau within the envelope was 0.08 ± 0.06 (mean ± s.d., n=50 envelopes in 10 independent experiments), and outside the envelope 0.12 ± 0.10 (n=50 regions in 10 independent experiments). Two-sided t-test, p=6.63*10^-4^. **e.** Density of tau within the envelope region for phospho-tau (magenta) and dephosho-tau (cyan) envelopes. Tau density in phospho-tau envelopes: 63.0 ± 13.5 (0 – 10% coverage), 72.1 ± 19.4 (10-20%), 72.4 ± 10.7 (20-30%), and 87.2 ± 95.6 (30-50%) (mean ± s.d., n=18, 16, 18, 12 envelopes in 5, 3, 4, 1 independent experiments); in dephospho-tau envelopes: 78.3 ± 15.0 5 (0 – 10% coverage), 79.3 ± 16.0 (10-20%), 83.3 ± 16.5 (20-30%), 108.2 ± 28.8 (30-50%) (n=7, 17, 30, 37 envelopes in 2, 2, 4, 5 independent experiments. Two-sided t-test between tau samples (left to right): p=0.0207, p=0.2569, p=0.0165, p=0.0169. **f.** Envelope disassembly rate after addition of active Cdk5 (magenta), deactivated Cdk5 (cyan), or in absence of Cdk5 (control, grey). Envelope disassembly rate for active Cdk5: 16.4 ± 13.7 nm/s (mean ± s.d., n=99 envelopes in 7 independent experiments); for deactivated Cdk5 3.2 ± 3.8 nm/s (n=123 envelopes in 8 independent experiments); for control 1.5 ± 1.1 nm/s (n=38 envelopes in 4 independent experiments). Two-sided t-test (left to right): p=2.30*10^-20^, p=5.92*10^-10^, p=0.0075. **g.** Fluorescence micrographs of removal of 10 nM Bact-tau in presence of active Cdk5 (left panels, Bact-tau in magenta) or deactivated Cdk5 (right panels, Bact-tau in cyan). Microtubules (black) are visualized using IRM. Envelope disassembly from the boundaries is indicated by the white arrows while fission events within the boundaries of the envelopes are indicated by yellow arrows (only observable in the active Cdk5 example). Note the different timepoints and intervals for the two samples. Scale bars: 2 μm. **h.** Number of fission events within the boundaries of Bact-tau envelopes during disassembly in presence of Cdk5 (magenta), deactivated Cdk5 (cyan), or in absence of kinase (control, grey). Number of envelope fissures in presence of active Cdk5: 0.75 ± 1.02 mm^-1^s^-1^ (mean ± s.d., n=117 envelopes in 7 independent experiments); deactivated Cdk5: 0.03 ± 0.17 mm^-1^s^-1^ (n=119 envelopes in 8 independent experiments); in absence of kinase (control): 0.16 ± 0.34 mm^-1^s^-1^ (n=52 envelopes in 4 independent experiments). Two-sided t-test (left to right): p=8.85*10^-13^, p=7.39*10^-5^, p=0.001364.

Knowing that phosphorylated tau participates in envelope formation, we hypothesized that the lowered propensity to form envelopes is caused by a combination of (i) a reduced interaction of phosphorylated tau with the microtubule lattice, as observed previously^11,12,19–21^, and (ii) reduced cohesiveness of the envelope, presumably due to reduced tau-tau interaction between phosphorylated tau molecules. To demonstrate (i) the reduction of the affinity of phosphorylated tau molecules on microtubules, we investigated the density of tau on the surface of microtubules. We performed the tau-microtubule interaction experiment at two different conditions; on GMPCPP-microtubules (which prevent the envelope formation) and on native GDP-microtubules (where tau binds preferentially in the envelope form^15^) (Methods, Supplementary Fig. 2c,d,e). Consistent with previous findings^11,12,19–21^, at most conditions, except for the saturating conditions on GDP-lattices, we found that phospho-tau is present on the microtubule lattice at lower densities compared to dephospho-tau (Supplementary Fig. 2c,d,e), confirming that phosphorylation reduces tau-microtubule interaction. To test if, furthermore, phosphorylation of tau, (ii) influences the cohesiveness of the envelopes, we mixed 8 nM mCherry-labeled phospho-tau with 8 nM GFP-labeled dephosho-tau, added this mixture to surface immobilized microtubules and observed the formation of envelopes. In line with the results of our pelleting assay, we observed that these envelopes exhibited both GFP and mCherry fluorescence (Fig. 2c), demonstrating that phosphorylated tau molecules are competent to participate in envelope formation. We then analyzed the ratio of phospho-tau to dephospho-tau outside of the envelope region and compared it to the ratio within the envelopes. This analysis revealed a significant relative decrease of phospho-tau within the tau envelopes compared to the regions outside the envelopes (Fig. 2d), showing that, additional to lower affinity to the microtubule lattice, phosphorylated tau less readily participates in envelope formation. Next, we prepared tau envelopes of either phospho- or dephospho-tau and studied the density of tau molecules within the enveloped regions. We found that for any given envelope coverage, the density of tau within the envelope region is lower in envelopes prepared from phospho-tau compared to envelopes prepared from dephospho-tau (Fig. 2e). This data suggests that envelopes formed by phosphorylated tau consist of a less dense and potentially more gap-prone structure. To test this hypothesis, we prepared tau envelopes on surface-immobilized microtubules using 10 nM Bact-tau, that has a low phosphorylation degree, and removed tau from solution to observe the disassembly of the envelopes. The removal of tau from solution was either performed in presence of (i) active Cdk5 kinase, (ii) deactivated Cdk5 kinase, or (iii) in the absence of a kinase (control) (Methods). In presence of active Cdk5, we observed that the disassembly of the envelopes was significantly faster compared to the conditions with deactivated Cdk5 or the control (Fig. 2f,g(note the different experimental timeframes), Supplementary Movie 5,6). These findings indicate that the Cdk5 kinase actively phosphorylated the tau in the envelopes, causing envelope destabilization. Interestingly, we observed a striking increase in the number of fission events within the envelopes in the presence of active Cdk5, compared to the deactivated Cdk5 or control envelopes (Fig. 2g,h). Combined, these data suggest that tau phosphorylation compromises the integrity and cohesiveness of the tau envelopes.

### Tau phosphorylation affects envelope formation in cells

We next investigated how envelope formation is regulated by tau phosphorylation in a cellular environment. Initially, we performed TIRF assays with cell lysates (Supplementary Fig. 3a, Methods). To generate tau lysates, we either overexpressed GFP-tau in HEK cells (denoted as HEK lysate), or we overexpressed GFP-tau together with proline-directed kinase Cdk5 and its activator p25 (Cdk5/p25) in HEK cells (denoted as Cdk5 lysate). We then lysed the cells, added the lysates to surface-immobilized microtubules, and followed the tau-microtubule interaction using TIRF microscopy (Supplementary Fig. 3a,b). While we observed tau envelopes forming in the HEK lysate, there were no envelopes forming in the Cdk5 lysate (Supplementary Fig. 3b,c). These observations are in line with our in vitro experiments showing that phosphorylation of tau decreases the propensity of tau to form envelopes. Next, we overexpressed GFP-tau and mScarlet-tubulin in U-2 OS cells and followed the tau signal correlated to the microtubule signal throughout the cell cycle, which is tightly regulated by activation of specific proline-directed cyclin-dependent kinases known to phosphorylate tau^22^. When following the tau signal throughout the cell cycle, we observed high tau signal on microtubules in interphase cells, where the overall activity of kinases is low, and a significant reduction of the tau signal on microtubules in mitotic cells (Supplementary Fig. 3d,e,f), where multiple mitotic kinases are active^23^. This observation is in agreement with our in vitro findings and published data of decreased affinity of phosphorylated tau for microtubules^11,12^.

To discern details of the tau-microtubule interaction in living cells and its regulation by phosphorylation, we overexpressed GFP-tau together with Cdk5/p25 in IMCD-3 cells, denoted as tau-Cdk5. In a control experiment, we overexpressed GFP-tau in IMCD-3 cells in the absence of Cdk5/p25, denoted as tau. Additionally, we overexpressed N-terminally truncated GFP-tau that is not able to form tau envelopes^13,14^, denoted as tau-ΔN (Methods). At elevated (micromolar) tau concentrations, tau envelopes are not readily discernable by a local increase in tau density^15^. We thus did not expect to observe regions of high and low tau density in cells, since tau is present at micromolar concentrations in cells^24,25^. Indeed, in both tau and tau-ΔN overexpressing cells, as well as in the tau-Cdk5 cells, tau covered microtubules uniformly along their entire lengths (Supplementary Fig. 3g). In tau-ΔN and tau-Cdk5 cells, tau signal on the microtubule was however weaker (Supplementary Fig. 3h), although the expression levels were comparable (Supplementary Fig. 3i), suggesting that in tau-ΔN and tau-Cdk5 cells, tau has a lower affinity for the microtubule surface or is bound at lower density because of reduced cohesiveness between the tau molecules.

To assess the turnover of tau on microtubules in our differently transfected cells, we used fluorescence recovery after photobleaching (FRAP). We photobleached a circular region of the cell that contained microtubules covered by tau, and studied the recovery of the tau signal on the microtubules over time (Fig. 3a, Supplementary Fig. 3j, Supplementary Movie 7-9). We found that the recovery of the fluorescent tau signal was slowest and the immobile fraction highest in our control cells expressing full length tau (Fig. 3b,c, Supplementary Fig. 3j), while in cells expressing tau-ΔN, which does not form envelopes, the recovery was fastest and there was no detectable immobile fraction. Since cooperatively bound tau shows lower turnover^14^, this data suggests that full length tau is bound to the microtubules in the form of a tau envelope. Interestingly, the recovery time and immobile fraction of the tau signal in tau-Cdk5 cells fell inbetween that of the control cells and the tau-ΔN cells, further supporting our in vitro data and indicating that phosphorylated tau is able to form envelopes, however, their turnover is faster and the envelopes are therefore less stable.

**Fig. 3.**
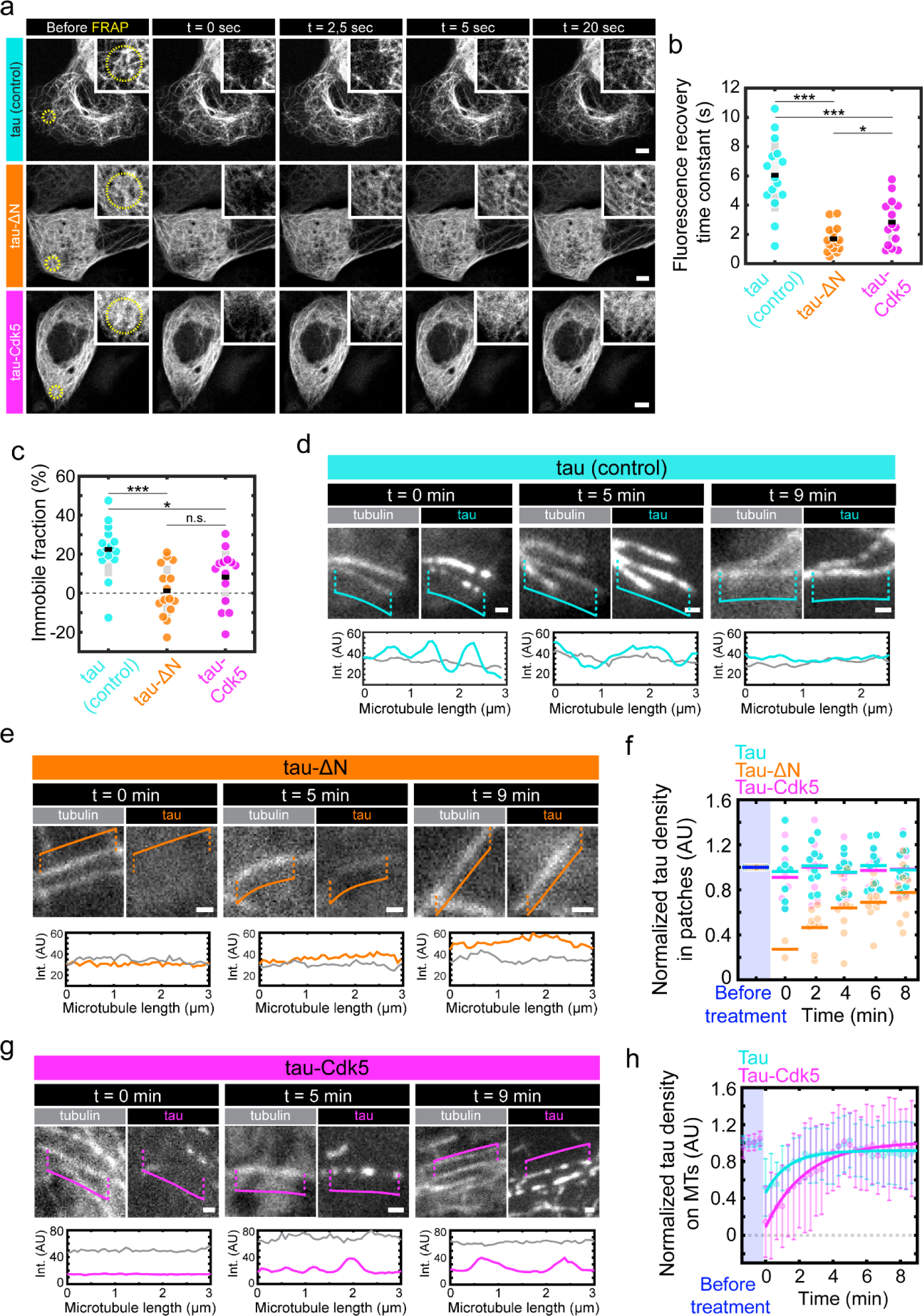
Tau phosphorylation affects envelope formation in cells. **a.** Fluorescence micrographs of FRAP experiment on IMCD-3 cells expressing GFP-tau (control, cyan), GFP-tau-ΔN (tau-ΔN, orange), or GFP-tau with Cdk5/p25 (tau-Cdk5, magenta) at different timepoints before and after FRAP. The FRAP-region is drawn as a yellow-dotted circle in the left panels (before FRAP), a zoom-in containing the FRAP-region is shown in the top right corner of each micrograph. Scale bars: 5µm. **b.** Time constant of the fluorescence recovery of the tau signal in the different groups. Time constant in control cells was 6.05 ± 2.50 sec (mean ± s.d., n=15 cells in 15 independent experiments); in tau-ΔN cells was 1.69 ± 0.89 sec (mean ± s.d., n=15 cells in 15 independent experiments); in tau-Cdk5 cells was 2.83 ± 1.61 sec (mean ± s.d., n=14 cells in 14 independent experiments). Two-sided t-test p-values (from left to right): p=6.76*10^-7^, p=0.00034, p=0.0242. **c.** Immobile fraction measured from fluorescence recovery curve. Immobile fraction in control cells was 22.49 ± 13.77 % (mean ± s.d., n=15 cells in 15 independent experiments); in tau-ΔN cells 1.14 ± 13.15 % (mean ± s.d., n=15 cells in 15 independent experiments); in tau-Cdk5 cells 8.23 ± 14.52 % (mean ± s.d., n=14 cells in 14 independent experiments). Two-sided t-test p-values (left to right): p=0.000167, p=0.0114, p=0.179. Fluorescence micrographs of IMCD-3 cells expressing mScarlet-tubulin and GFP-tau in control cells (tau, cyan) at t=0 min (left), t=5 min (middle), and t=9 min (right) after elevated-pH treatment. Corresponding linescans are shown below the micrographs. Different microtubules were selected at different timepoints due to dynamic behavior of the microtubules. Scale bars: 1µm. **e.** Fluorescence micrographs of mScarlet-tubulin and GFP-tau signal in tau-ΔN cells (orange) at t=0 min (left), t=5 min (middle), and t=9 min (right) after elevated-pH treatment. Corresponding linescans are shown below the micrographs. Different microtubules were selected at different timepoints due to dynamic behavior of the microtubules. Scale bars: 1µm. **f.** Normalized tau density in patches in GFP-tau control cells (tau, cyan) and GFP-tau-Cdk5/p25 cells (tau-Cdk5, magenta). Normalized tau density along the microtubule lattice (no patches are visible) in GFP-tau-ΔN cells (tau-ΔN, orange). Tau densities were measured at 5 timepoints after elevated-pH treatment and normalized to the tau density along the microtubules before the treatment (dark blue). **g.** Fluorescence micrographs of mScarlet-tubulin and GFP-tau signal in tau-Cdk5 (magenta) at t=0 min (left), t=5 min (middle), and t=9 min (right) after elevated-pH treatment. Corresponding linescans are shown below the micrographs. Different microtubules were selected at different timepoints due to dynamic behavior of the microtubules. Scale bars: 1µm. **h.** Time-trace of tau density on the MTs in the whole cell after elevated-pH treatment normalized to the tau density on the MT before the treatment in GFP-tau cells (tau, cyan) and GFP-tau-Cdk5/p25 cells (tau-Cdk5, magenta). Exponential time constant was 1.3 min for tau cells and 2.5 min for tau-Cdk5 cells (Methods).

Tau binding is sensitive to pH of the environment in vitro as well as in cells^26,27^. It was shown that changing the pH of the media of cells manifests in the change of the intracellular pH, which can rapidly affect binding of tau to microtubules in cells^26^. As the presence of envelope-incorporated tau can be detected during the assembly or disassembly of the envelopes^14,15^, we used pH change to directly test the presence of tau envelopes in cells by exchanging the media during imaging for a solution with slightly higher pH (from pH 7.4 to pH 8.4, Methods). In these conditions, any tau molecules that are non-cooperatively bound to the microtubule, due to their much higher turnover compared with tau in the envelopes, would be readily released from microtubules, while cooperatively bound tau forming envelopes would be more resilient to disassembly. Indeed, in our positive control cells expressing full length GFP-tau (denoted as tau), we observed gaps forming in the tau signal (manifested as increased coefficient of variation of the tau signal along the microtubule, Supplementary Fig. 3g,k) while the microtubules remained unaffected (coefficient of variation of the tubulin signal is not affected, Supplementary Fig. 3g,k). These gaps in the tau signal left clearly separated tau patches on the microtubules, strongly suggesting the presence of cooperatively binding tau molecules forming envelopes, constituting the observed patches (Fig 3d, t=0 min), and confirming our previous observations, where taxol treatment led to dissociation of tau from microtubules in cells in a patch-like pattern^15^. The advantage of the elevated pH treatment compared to the taxol treatment is that the tau unbinding is faster and reversible, which allowed us to analyze the re-binding of tau to microtubules. Presumably due to a recovery of the intracellular pH after the treatment, the gaps started to close and the tau patches regrew (Fig 3d, t = 5 and 9 min), which is analogous to the growth of envelopes in vitro, when excess tau is available^14,15^. During the whole process, the density of the GFP-tau signal (GFP intensity per unit length) in the patches remained constant (Fig. 3f), further suggesting that these patches represent tau envelopes (which grow and shrink only at the boundaries). Within 9 minutes after the elevated-pH treatment, the microtubules regained full tau coverage (Fig. 3d, Supplementary Fig. 3g (whole cell images),k,l, Supplementary Movie 10). When we performed the elevated-pH treatment on tau-ΔN (incompetent of envelope formation), we did not detect any tau patches on the microtubules after the elevated-pH treatment (Fig 3e, t=0 min). Instead, we observed that the tau signal was removed uniformly and fully along the entire lengths of the microtubules (Fig. 3e, Supplementary Fig. 3g (whole cell images),k,l, Supplementary Movie 11), which suggests that tau molecules were bound to the microtubules individually, non-cooperatively. Subsequently, when following the recovery after elevated-pH treatment, we observed that the tau-ΔN signal returned, again uniformly, along the microtubule lattice. Unlike full-length tau molecules, which exhibited regrowth of the patches while remaining a constant density within the patches, the density of tau-ΔN signal on the microtubule uniformly increased during recovery (Fig. 3e,f, Supplementary Fig. 3g (whole cell images),k,l, Supplementary Movie 11). This further suggests that ΔN-tau, in these cells, does not form envelopes, and instead binds to microtubules non-cooperatively (i.e. with high turnover and high diffusivity). Strikingly, when following the recovery after elevated-pH treatment in the tau-Cdk5 cells, we observed that the reappearance and recovery of the signal occurred in a patch-like manner, closely resembling the reappearance of the tau signal in our positive control cells (Fig 3g, Supplementary Fig. 3g (whole cell images),k,l, Supplementary movie 12). Moreover, the density of the tau signal on microtubules within the patches remained constant throughout the treatment in the tau-Cdk5 cells (Fig. 3f), indicating that tau was bound cooperatively to the microtubules. Interestingly, the reappearance of the tau signal on microtubules was slower in tau-Cdk5 cells than in the control cells indicating that tau phosphorylation leads to slower envelope recovery (Fig. 3h). Combined, these data suggest that in control and tau-Cdk5 cells, the patches and the high-density tau areas covering microtubules consisted of cooperatively bound tau molecules forming cohesive tau envelopes, while N-terminally truncated tau in tau-ΔN cells bound non-cooperatively only. The fact that after the elevated-pH treatment, tau signal disappeared almost completely and reappeared slower in the tau-Cdk5 cells compared to cells overexpressing control tau is in agreement with our in vitro data showing that tau phosphorylation does not prevent tau envelope formation but makes tau less prone to form envelopes and makes the resulting envelopes less stable. Combined, our data suggests that tau phosphorylation decreases the cohesiveness of tau envelopes, thereby negatively affecting the stability of the tau envelopes in living cells.

### Tau phosphorylation affects envelope functionality in vitro and in living cells

Finding that tau envelopes could be formed by tau in both its phosphorylated and non-phosphorylated states, albeit at different efficiencies, and that phosphorylation leads to compromised envelope integrity, we asked if tau phosphorylation additionally affects the functionality of the envelope. Tau envelopes regulate the accessibility of the microtubule surface for other microtubule-associated proteins, such as molecular motors or microtubule severing enzymes, thereby regulating their function^13,14^. To test if envelopes formed by phospho-tau and dephospho-tau differentially regulate typical molecular motors, like kinesin-1, and typical severing enzymes, like katanin, we employed in vitro reconstitution: in separate measurement chambers, we formed envelopes of similar sizes either with dephospho-tau (5 nM) or phospho-tau (30 nM). Keeping the tau concentrations in solution, we then added kinesin-1 to the measurement chamber and studied the effect of the tau envelopes on kinesin-1. In accordance with previous findings^14^, we found that molecules of kinesin-1 could walk on microtubules outside of the envelope regions and were mostly excluded from the envelopes (Supplementary Fig. 4a, Supplementary Movie 13,14). Quantifying the landing rate of kinesin-1 inside the envelope regions, we found no significant difference between the functioning of dephospho-tau and phospho-tau envelopes (Supplementary Fig. 4b). We then repeated the experiment with microtubule severing enzyme katanin. In line with previously published data^14,28^, we observed that the regions of microtubules, which were covered only by diffusible tau and not by tau envelopes, were quickly disintegrated by katanin (Fig. 4a), confirming that the cooperatively-bound tau and not the diffusibly-bound tau, can protect microtubules against katanin severing. The envelope-coated regions of the microtubule were protected and prevailed for longer periods. While the density of tau envelopes remained constant (Supplementary Fig. 4c), the envelopes were slowly disassembled by katanin from their boundaries, with occasional severing events within the enveloped region (Fig. 4a, Supplementary Movie 15). The disassembly rate of enveloped regions from their boundaries did not show any significant difference between phospho-tau and dephospho-tau envelopes (Fig. 4b). However, we observed significantly increased rates of katanin severing events within the envelope regions formed by phosphorylated tau compared to non-phosphorylated tau (Fig. 4c), which may be explained by the reduced density of tau within envelopes formed by phospho-tau and reduced tau envelope integrity, as shown above. Combined, these experiments show that envelopes constituted by either phosphorylated or non-phosphorylated tau similarly inhibit kinesin-1 movement, while differentially protect against katanin-mediated severing. Tau phosphorylation thus, in addition to reducing the propensity of tau molecules to bind cooperatively, furthermore reduces the protective functionality of the formed tau envelopes.

**Fig 4.**
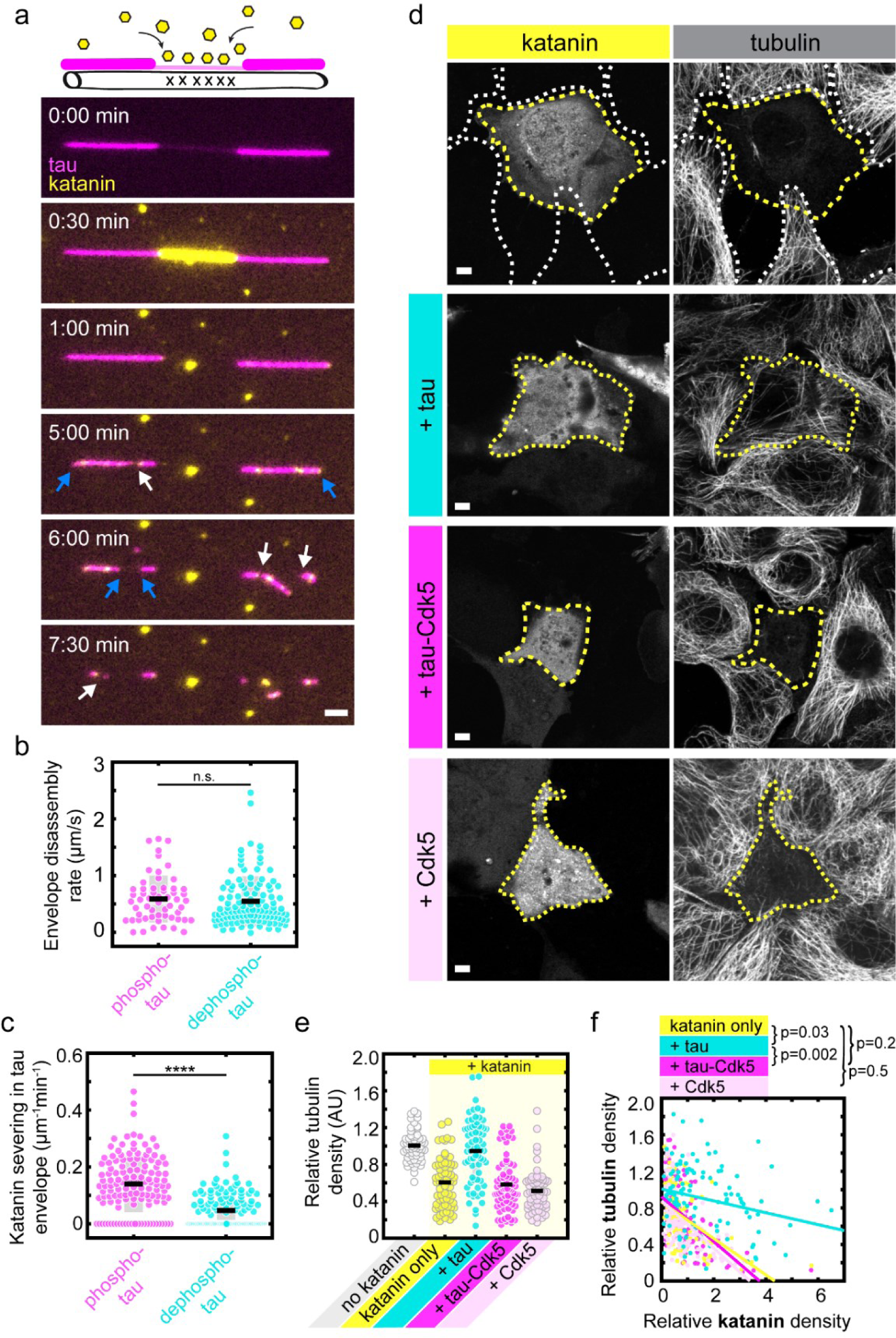
Tau phosphorylation affects envelope functionality in vitro and in living cells. **a**. Schematics and multichannel fluorescence micrographs showing rapid katanin-GFP-mediated severing of a microtubule region not covered by a tau envelope (panel 2, tau in magenta, katanin in yellow) and subsequently the much slower disassembly of the microtubule regions protected by a tau envelope (panels 3-6). Katanin severing leading to disassembly of envelope-covered portions of the microtubule from their boundaries is indicated by blue arrows, while occasional katanin severing within a tau envelope (severing event) is indicated by white arrows. Scale bar: 2 μm. **b**. Envelope disassembly rate from the boundaries due to katanin severing of phospho-tau envelopes (magenta) was 0.59 ± 0.41 µm/s; dephospho-tau envelopes (cyan) was 0.50 ± 0.50 µm/s (mean ± s.d., n=57, 112 envelopes in 4,3 experiments). Two-sided t-test, p=0.2638. **c**. Katanin severing events within tau envelope boundaries in phospho-tau envelopes (magenta) was 0.14 ± 0.10 µm^-1^min^-1^; dephospho-tau envelopes (cyan) was 0.05 ± 0.06 µm^-1^min^-1^ (mean ± s.d., n=127,132 envelopes in 4,4 experiments). Two-sided t-test, p=1.85*10^-17^. **d.** Fluorescence micrographs of IMCD-3 cells expressing katanin-GFP (left panels) and stained to visualize tubulin (right panels). Cells were fixed and stained 12 hours after transfection. Additionally, cells are expressing tau (+ tau, cyan), tau and Cdk5/p25 (+ tau-Cdk5, magenta), or Cdk5/p25 in the absence of tau (+ Cdk5, light pink). Cells expressing katanin are marked with a yellow dotted line and cells not expressing katanin (non-transfected cells) are marked with white dotted lines. Scale bars: 5µm. **e.** Relative tubulin density 12 hours after transfection. Relative tubulin density in ‘no katanin’ cells was 1.01 ± 0.16 (white), in ‘katanin-only’ cells was 0.61 ± 0.27 (yellow), in ‘+ tau’ cells was 0.95 ± 0.35 (cyan), in ‘+ tau-Cdk5’ cells was 0.58 ± 0.28 (magenta), in ‘+ Cdk5’ cells was 0.51 ± 0.23 (light pink) (mean ± s.d., n=60, 64, 68, 71, 60 cells in 3 independent experiments). **f.** Correlation of the relative tubulin density (y-axis) compared to the relative katanin density (x-axis). Correlation coefficients of ‘katanin only’ is −0.49 (yellow), of ‘+ tau’ is −0.25 (cyan), of ‘+ tau-Cdk5’ is −0.57 (magenta), and of ‘+ Cdk5’ is −0.54 (light pink). Correlation coefficient comparisons are indicated above the graph.

We next asked if tau phosphorylation affects the functionality of envelopes in living cells. To test this, we prepared IMCD-3 cells overexpressing katanin-GFP (denoted as ‘katanin only’), or IMCD-3 cells overexpressing katanin-GFP in combination with mCherry-tau (denoted as ‘+ tau’), or mCherry-tau and Cdk5/p25 (denoted as ‘+ tau-Cdk5’). As a control we prepared IMCD-3 cells overexpressing katanin-GFP in combination with Cdk5/p25 without overexpression of mCherry-tau (denoted as ‘+ Cdk5’). Cells were fixed 12 hours after transfection and stained for tubulin to visualize the presence of microtubules (Methods). In cells expressing katanin in absence of tau (Fig. 4d, transfected cells, yellow marker), a clear reduction in tubulin signal was observed in comparison with cells not expressing katanin (Fig. 4d,e, non-transfected cells, white marker). Consistent with previously published data^6,14^, in cells expressing katanin in combination with tau, no reduction in tubulin signal was measured, indicating that tau protected microtubules against the severing activity of katanin, which is in agreement with our in vitro data. Strikingly, phosphorylation of tau by overexpression of Cdk5/p25, impeded this protective ability, resulting in a reduction in tubulin signal to similar level as in cells expressing katanin only or in cells expressing katanin in combination with Cdk5/p25 (Fig. 4d,e). Comparing the relative density of tubulin with the relative density of katanin 12 hours after transfection, we found that tubulin density inversely correlated with katanin density in all cell groups (Fig. 4f). This correlation was weaker in presence of tau, and stronger in presence of tau-Cdk5, further supporting the notion that phosphorylation of tau decreases its protective functionality (Fig. 4f). Combined, these experiments show that phosphorylation of tau impedes its protective functionality both in vitro and in living cells.

## Discussion

Tau has been shown to modulate many neurodevelopmental processes through its interaction with microtubules, while its deregulation is associated with pathogenesis of numerous neurodegenerative disorders. It is known that phosphorylation regulates tau, but the detailed mechanism of how it affects tau function is still ill-understood. Here we show that phosphorylation of tau controls the formation of the protective tau envelopes. Tau envelopes are cohesive patches of cooperatively binding tau molecules characterized by limited lateral translocation and low turnover of the constituting tau molecules, which enclose the microtubule lattice and are selectively permeable for some proteins, while protecting the microtubules from others, for example the severing enzyme katanin^14^. This mode of tau binding to microtubules is contrasted by non-cooperative interactions of tau molecules binding individually, observed in vitro at low tau concentration^13,14^ and upon disturbance of the cellular homeostasis, for example by pH-shock (Fig.4d-g). This non-cooperative mode of tau binding is characterized by rapid diffusion and high turnover of the individual tau molecules, which, in absence of cohesion, does not shield against the binding of other proteins, and thus leaves the microtubule lattice vulnerable to severing enzymes like katanin^14^. Here, we showed that phosphorylation of tau impedes the envelope formation and integrity and reduces their protective functionality.

Previous results showed that tau phosphorylation reduces tau association with microtubules ^19–21,29^. In accordance with these findings, we observed, outside of the tau envelopes, reduced binding of phosphorylated tau to microtubules, compared to non-phoshorylated tau. We suggest that increased binding of non-phosphorylated tau to the microtubule lattice and consequent higher tau densities on the microtubule lead to an increase in frequency of tau-tau encounters on the microtubule lattice, leading to a higher probability of nucleation of a tau envelope and increased envelope growth. Additionally, our finding that the relevant phosphorylation sites are located in the proline-rich regions and the C-terminus of tau (Fig. 2b), but not in the microtubule binding repeats (a) demonstrates that phosphorylation of tau domains other than the binding repeats play a significant role in the interaction of tau with the microtubule lattice, and (b) suggests that tau-tau interaction might play a significant role in the envelope formation. This notion is consistent with the observation that tau molecules, under certain conditions, can phase-separate^30^ and that phosphorylation regulates condensate formation^31^. For both reasons (a) and (b), the concentration required to achieve comparable microtubule envelope coverage, is much higher for phosphorylated tau as compared to non-phosphorylated tau. At any given tau concentration, phosphorylated tau generated envelopes which cover, and thus protect, a smaller fraction of the microtubule surface, leaving it completely uncovered at low nanomolar concentrations. The reduced protection resulting from tau phosphorylation could be due to higher turnover of phosphorylated tau, which might result in transient defects in the envelope. Thus, in cells, tau phosphorylation, at any given tau concentration, will dramatically reduce microtubule envelope coverage and thereby protection. And, importantly, due to the reduced protection, microtubule instability can be observed even before tau envelope disassembly occurs.

As non-phosphorylated tau more readily participates in envelope formation, our data suggests that the different affinities are not the only reason for the observed effect of the phosphorylation on tau envelope behavior. Our data suggest that, additionally, the increased envelope coverage observed for non-phosphorylated tau may be due to stronger tau-tau interactions on the microtubule surface, creating a more cohesive and impenetrable structure. Tau phosphorylation weakens these interactions and creates a structure that is easier to penetrate by microtubule severing enzymes such as katanin. This notion is corroborated by the finding that phosphorylated tau forms envelopes that have lower tau density compared to dephosphorylated tau envelopes (suggesting weaker tau-tau interactions) and that katanin severs microtubules even within the boundaries of envelopes consisting of phosphorylated tau. This hypothesis is especially plausible when considering the position of phosphorylation in our phospho-tau samples: While little phosphorylation was detected within the binding repeats (R1-R4 and R’), which directly interact with the microtubule, most phosphorylation was detected in the projection domains of tau (in particular P1, P2, and the C-terminus), which are the regions thought of as establishing tau-tau interaction^13,14^.

Increased tau phosphorylation has been detected in multiple neurodegenerative disorders and has been associated with formation of phosphorylated tau aggregates^32,33^. The specific effect of tau phosphorylation in the pathogenesis of different diseases is still being analyzed. Using in vitro and in vivo models we demonstrate that increased phosphorylation interferes with formation of protective tau envelopes. Moreover, we show that the presence of phosphorylated tau leads to disassembly of already formed tau envelopes. This can critically amplify the deleterious effect of phospho-tau on neurons as release of tau from envelopes may locally increase its concentration and promote formation of phosphorylated tau aggregates with additional toxic effect on neuronal transport and function.

Tau is phosphorylated by numerous kinases in healthy neurons, but deregulation of Cdk5 seems critical in the neuropathological process leading to neurodegeneration. We now demonstrate that upregulation of Cdk5 has a deleterious effect on microtubule protective functions of tau envelope formation and maintenance, and that microtubules covered by tau envelopes formed by phosphorylated tau are more prone to disintegration, e.g. by microtubule severing enzymes, such as katanin. Thus, by linking Cdk5 upregulation with tau phosphorylation and tau envelope disintegration, our work provides a mechanism of microtubule regulation in cells and in pathogenesis of neurodegenerative disorders, such as Alzheimer’s disease. Neurodegeneration and microtubule destabilization is often linked to tau hyper-phosphorylation. Our results suggest that microtubule destabilization could be the result of the impaired protective functionality of tau envelopes upon phosphorylation of tau.

## Methods

### Protein constructs and purification

#### Insect cell expressed tau

For in vitro experiments, GFP- or mCherry-labelled tau (h441-tau; NM_005910.6:151-1476) was expressed in insect cell and purified using the baculovirus expression system (DefBac DNA). Sf9 cells were infected with 8 ml of P2 baculovirus stock (1:100 ratio of P2 virus to cell culture), incubated at 27°C with moderate shaking, and harvested 72 hours post infection. Cells were harvested by centrifugation at 300 x g for 10 min and resuspended in PBS before snap-freezing the cells, or prior to purification in lysis buffer (25 mM HEPES pH 7.4, 150 mM KCl, 20 mM imidazole, with 1 mM DTT, benzonase (1.25 µL of 25 U/µl, 70664, Novagen) and 1x Protease inhibitor cocktail (34044100, Roche Diagnostics GmBH). Cells were lysed by spinning at 70000 x g for 1 hour at 4°C and collecting the supernatant. The lysate was incubated with NiNTA agarose resin (XF340049, Thermo Scientific) HiTrap for 2 hours at 4°C by slowly rotating. After incubation, beads were washed with 3×20 ml wash buffer (25 mM HEPES, 150 mM KCl (or 700 mM KCl in wash step 2), 1mM DTT, 20 mM imidazole). 6xhis tag was removed by incubating the beads with PreScission protease (homemade 3C HRV protease, 1:100, 1µg enzyme/100 µg of protein, overnight at 4°C while rotating). The next day the cleaved protein was collected and concentrated by spinning the sample at 3500 RPM at 4°C using −50kDa centrifugal filter tube (Amicon Ultra-15, Merck). The protein was purified by size-exclusion chromatography using a Superdex 200 10/300 GL column (GE28-9909-44, Sigma) with an NGC Chromatography system (Bio-Rad), equipped with ChromLab software (Bio-Rad) in 25 mM HEPES pH 7.4, 150 mM KCl, 1 mM DTT, 0.1 mM ATP, 1 mM EDTA. Collected peak fractions were concentrated to 10-40 µM using −50kDa centrifugal filter tube (Amicon Ultra-15, Merck). Protein concentration was measured with a NanoDrop ND-1000 spectrophotometer (Thermo Scientific) at 280 nm absorbance. Proteins were flash-frozen in liquid nitrogen and stored at −80°C. All steps in the purification were performed at 4°C.

#### Bacterial cell expressed tau

Fluorescently tagged tau used for in vitro experiments (h441-tau was subcloned into the expression vector based on pET11Kan-N-HIS6-3C-mNeonGreen or pET11Kan-N-HIS6-3C-mRuby3) was expressed in E. coli BL21(DE3)-RIPL strain. The cells were grown at 30°C until OD_600_ of 0.5-0.6, the protein expression was then induced by 0.1 mM IPTG, and the cells were grown overnight at 16°C. Bacterial cells (3-4 g) were lysed in 45 ml of lysis buffer (50 mM Tris pH 8.0, 300 mM NaCl, 2 mM bME, 20 mM Imidazole, 0.5 µL Benzonase, and 1x Protease inhibitor cocktail), sonicated (5 min; On/Off: 2/4 s) and centrifuged (40000 x g; 30 min; 4 °C). The soluble fraction was then subjected to Strep-Tactin XT purification (washing buffer: 50 mM Tris pH 8.0, 300 mM NaCl, 2 mM bME), and eluted using BXT buffer (100 mM Tris pH 8.0, 150 mM NaCl, 1 mM EDTA, 50 mM Biotin). The purified protein was concentrated using VivaSpin-10kDa-HY and subjected to Size exclusion chromatography (see Insect cell expressed tau; buffer: 50 mM Tris pH 8.0, 300 mM NaCl, 1 mM DTT). Protein concentration was measured with a NanoDrop (see Insect cell expressed tau) at 280 nm absorbance. Proteins were flash-frozen in liquid nitrogen and stored at −80°C. All purification steps were performed at 4°C.

#### Katanin + Kinesin expression and purification

Katanin-GFP^34^ (p60 & p80-GFP)and kinesin-1-GFP^35^ were expressed and purified as previously described.

### TIRF microscopy

Total internal reflection fluorescent (TIRF) microscopy experiments were performed on an inverted microscope (Nikon-Ti E, Nikon TI2 E) with H-TIRF module or iLas2 equipped with 60x or 100x NA 1.49 oil immersion objectives (Apo TIRF or SR Apo TIRF, respectively, Nikon) and CMOS Hamamatsu Orca Flash 4.0 LT, sCMOS Hamamatsu ORCA 4.0 V2, or PRIME BSI (Hamamatsu Photonics, Teledyne Photometrics) cameras. Microtubules were visualized using interference reflection microscopy (IRM) and fluorescent proteins by switching between microscope filter cubes for EGFP, mCherry, and Cy5 channels or by using a quad band set (405/488/561/640). The microscopes were controlled with Nikon NIS Elements software (v5.02, v5.20 or v5.42). All experiments were performed at room temperature by several experimentalists over the course of multiple months. No data was excluded from the study.

#### Experimental chamber preparation

For TIRF experiments, chambers were assembled by melting thin strips of parafilm in between two glass coverslips silanized with 0.05% dichlorodimethylsilane (DDS, #440272, Sigma). The chambers were incubated with 20 µg/mL anti-biotin antibodies (in PBS, #B3640, Sigma) for 5 min or 20 µg/mL anti-β-tubulin antibodies (in PBS, #T7816, Sigma) for 5 min, followed by 1% Pluronic (F127 in PBS, #P2443, Sigma) for at least 30 min. (Biotin-labeled) Microtubules (Methods) were diluted into BRB80T (BRB80: 80mM PIPES pH 6.9, 1mM EGTA, 1mM MgCl2, supplemented with 10 µM paclitaxel (#17191, Sigma)), then incubated in the chamber and allowed to adhere to the antibodies for 30 sec. Unbound microtubules were washed away with BRB80T and chambers were pre-incubated with TIRF assay buffer AB (50 mM HEPES pH 7.4, 1 mM EGTA, 2 mM MgCl_2,_ 75 mM KCl, 10 mM dithiothreitol, 0.02 mg/ml casein, 10 µM taxol, 1 mM Mg-ATP, 20 mM D-glucose, 0.22 mg/ml glucose oxidase and 20 µg/ml catalase) prior to experiments. Unless stated otherwise, all experiments were conducted in TIRF assay buffer (AB). All experiments were quantified by pooling data from multiple chambers performed on at least two different days. Chambers were never re-used for additional experiments.

#### Microtubule Assembly

Porcine brains were obtained from a local abattoir and used within ∼4 h of death. Porcine brain tubulin was isolated using the high-molarity PIPES procedure^36^. Biotin-labeled tubulin was purchased from Cytoskeleton Inc. (#T333P) and diluted 1:50 with unlabeled porcine brain tubulin to obtain biotin-labeled tubulin mix for surface-immobilization assays using biotin antibodies.

Taxol-stabilized microtubules (GTP polymerized, then taxol-stabilized; stored and imaged in presence of taxol) were polymerized from 4 mg/ml tubulin for 30 min at 37°C in BRB80 supplemented with 4 mM MgCl_2_, 5% DMSO, and 1mM GTP (#NU-1012, Jena Bioscience). The polymerized microtubules were diluted in BRB80T and centrifuged for 30 min at 18000 x g in a Microfuge 18 Centrifuge (Beckman Coulter). After centrifugation the pellet was resuspended and kept in BRB80T at room temperature.

GMPCPP-microtubules (GMPCPP polymerized, then taxol-stabilized; stored and imaged in presence of taxol) were polymerized from 4 mg/ml tubulin for 2 h at 37°C in BRB80 supplemented with 1mM MgCl_2_ and 1mM GMPCPP (#NU-405, Jena Bioscience). The polymerized microtubules were centrifuged for 30 min at 18000 x g in a Microfuge 18 Centrifuge (Beckman Coulter). After centrifugation the pellet was resuspended and kept in BRB80T at room temperature.

GDP-microtubules (GTP polymerized, then glycerol-stabilized; stored and imaged in presence of 40% glycerol) were polymerized as described for taxol-stabilized microtubules. After polymerization, the microtubules were gently diluted in BRB80-Gly40 buffer (80 nM PIPES, 2 mM MgCl_2_ and 1 mM EGTA, pH 6.8, 40% glycerol) and centrifuged as described above. After centrifugation the supernatant was discarded, and the pellet was resuspended gently in 50 μl of BRB80-Gly40. Microtubules were then kept at room temperate at least 1 hour (maximum overnight) before usage.

#### Tau sample preparation

To study the effect of phosphorylation of tau on envelope formation 4 samples were produced with various degrees of phosphorylation, as described below:

##### Phospho-tau

Tau expressed in insect cells, treated with buffer in absence of Alkaline Phosphatase. 2μM (0,2 mg/ml) insect cell expressed tau was incubated in 1x Fast Phosphatase Buffer (stock 10x) for 15 min at 37°C.

##### Dephospho-tau

Tau expressed in insect cells, treated with Alkaline Phosphatase (FastAP Phosphatase, #EF0651, Themofisher). 2μM (0,2 mg/ml) insect cell expressed tau was incubated with 2,5 g/mol Alkaline Phosphatase (stock 10 g/mol) and 1x Fast Phosphatase Buffer (stock 10x) for 15 min at 37°C.

##### Bact-tau

Tau expressed in bacterial cells, treated with buffer in absence of Cdk5/p35 kinase. 2μM (0,2 mg/ml) Bact-tau was incubated in Reaction Buffer A (K03-09, stock 5x) supplemented with 50μM DTT, 50μM ATP for 15 min at 37°C.

##### Bact-Cdk5-tau

Tau expressed in bacterial cells, treated with Cdk5/p35 kinase (#V3271, Promega). 2μM (0,2 mg/ml) Bact-tau was incubated in Reaction Buffer A (K03-09, stock 5x) supplemented with 50μM DTT, 50μM ATP and 0.02 μg/μl Cdk5/p35 kinase (stock 0.1 μg/μl) for 15 min at 37°C.

#### TIRF assays

In all TIRF experiments, chambers were prepared as described above and microtubules were observed using interference reflection microscopy (IRM). Movies were captured with appropriate frame interval and analysis was done after a certain incubation time as stated in the caption or methods.

##### Tau on microtubules

Tau samples were diluted in AB to the final concentration stated in the main text. After microtubule incubation, the diluted tau sample was added to the measurement chamber with at least four-fold amount of the chamber volume.

##### Cdk5 treatment in channel

Tau-mNeonGreen was diluted in AB buffer (supplemented with 0.5 mg/ml casein) to final concentration 15 nM and incubated on surface-immobilized microtubules for 10 min. After incubation, either i) active cdk5/p35 (activity: 0.1 μg/μl, diluted 10x) or ii) deactivated cdk5/p35 (deactivated by incubating at 95 °C for 10 min, diluted 10x), were added to the chamber, while tau concentration remained unchanged. Tau envelopes were observed for 15 min.

##### Kinesin-1 assay

Phospho-tau-mCherry or dephospho-tau-mCherry were diluted to concentrations at which envelopes of similar microtubule coverage assembled (30 nM and 5 nM, respectively). After microtubule incubation, tau was added to the measurement chamber and incubated for 3 minutes after which 25 nM kinesin-1-GFP was added in the presence of tau. Microtubules and the location of the tau envelopes were determined by taking a snapshot after which kinesin-1 was imaged for 30 sec using single molecule approach (no delay, 20 ms exposure time).

##### Katanin assay

Dephospho-tau-mCherry or phospho-tau-mCherry was diluted in AB and incubated on surface-immobilized microtubules for 5 min. Concentrations of dephospho- and phospho-tau were chosen such as to achieve similar microtubule coverage (0.8 nM for dephospho-tau and 3.5 nM for phospho-tau). After incubation, 100 nM katanin-GFP was added to the measurement chamber in presence of the established tau concentration. Chambers were imaged for 30 min with 5 sec interval.

##### Tau on GDP-microtubules

Tau-mCherry (insect cell expressed) was dephosphorylated using phosphatase as stated above. Dephosphorylated or phosphorylated (untreated) tau diluted in BRB80-Gly40-AB was added to surface immobilized GDP-microtubules (glycerol-stabilized) at concentrations: 0.2 nM, 1 nM, 2 nM, 4 nM, 10 nM, 50 nM or 100 nM, and incubated for 5 minutes. For the lower concentrations (0.2-4nM), tau was sequentially added to the same experimental chamber. For concentrations 10 nM and above, a new channel was used for every concentration. All experiments with Glycerol-stabilized microtubules were performed in BRB80-Gly40 buffer to ensure that microtubules remained stable. For in vitro TIRF assays, BRB80-Gly40 buffer was supplemented with 10 mM dithiothreitol, 20 mM d-Glucose, 1 mM ATP, 0.02 mg/ml Casein, 0.22 mg/ml Glucose Oxidase and 0.02 mg/ml Catalase (henceforth called BRB80-Gly40-AB).

##### Cdk5 treatment with tau envelope disassambly

Tau-mNeonGreen (bacterial expressed) was diluted in AB to a final concentration between 10-20 nM and incubated on surface-immobilized microtubules for 5 min. After incubation, tau was removed from the channel by the addition of 20 μl AB that contained either i) active Cdk5/p35 (activity: 0.1 μg/μl, diluted 10x), ii) deactivated Cdk5/p35 (deactivated by incubating at 95 °C for 10 min, diluted 10x) or iii) in absence of any kinase (i.e only AB, control). The disassembly of tau envelopes was observed for: 5 min (for active Cdk5); 10-45 min (for deactivated Cdk5); and 10 min (for control).

### TIRF Image analysis

Microscopy data were analyzed using ImageJ 2.3.0/1.53t (FIJI)^37^ and custom written Matlab (R2020b) codes. In images with substantial drift, the ‘StackregJ’ plugin was used to correct the drift (kindly provided by Jay Unruh at Stowers Institute for medical research in Kansas City, MO).

#### Kymographs

Kymographs were generated by drawing a line along the microtubule lattice and using the ImageJ kymographBuilder plugin.

#### Envelope coverage

Microtubule lengths were measured by using the IRM signal, tau envelope lengths were measured by using the fluorescent signal after 3 min of incubation. The envelope coverage represents the sum of all tau envelopes lengths divided by the sum of all microtubule lengths within one field of view.

#### Coverage difference after Cdk5 treatment

Tau envelope coverage difference was calculated by subtracting coverage by tau envelopes before the addition of active/deactivated cdk5/p35 and after 15 min after adding active/deactivated cdk/p35. Coverage was measured as described above.

#### Tau envelope growth rate

Length of the dephospho-tau envelopes was measured prior to (t=0 min) the 45 min incubation with either phospho- or dephospho-tau and then at the end of the experiment (t=45 min). The difference between the two lengths was calculated and subsequently divided by time to get the growing/shrinking rate (μm/min) of each tau envelope.

#### Tau density estimation

Tau density on the microtubules was measured in ImageJ by drawing a rectangle around the microtubule and measuring the mean. For background-subtraction the rectangle was then moved to an area directly adjacent to the microtubule where no microtubule is present and the mean was measured again and subtracted from the mean on the microtubule.

#### Affinities (Tau density plotted against tau concentration)

GMPCPP-lattice microtubules were immobilized on the coverslips surface and increasing concentrations of phosphorylated or dephosphorylated tau were added to the measurement chamber. The tau density was then measured on the microtubule lattice and plotted against the tau concentration in nM as a measure of the affinity of tau to the microtubule.

#### Fraction of phospho-tau in- or outside envelope region

Chambers were prepared as described above. 8.5 nM dephospho-tau-mCherry was mixed with 8.5 nM phospho-tau-mCherry (diluted in AB). The tau mixture was incubated on surface-immobilized microtubules for 5 min. The fraction of phospho-tau within the envelope was calculated as the density of phospho-tau in the enveloped region, divided by the total density of phospho- and dephospho-tau within the same region. The fraction of phospho-tau outside the envelope region was calculated as the density of phospho-tau in the non-enveloped region, divided by the total density of phospho- and dephospho-tau within the same region.

#### Envelope disassembly rate due to Cdk5 treatment

The envelope length was measured at the start of the video (before tau removal), and after 5-45 min (see above). Envelope lengths were determined by eye and measured using ImageJ. The disassembly rate was calculated as the difference in the envelope lengths at the beginning and at the end of the video, divided by the time between the two measurements. If an envelope disappeared fully before the end of the video, the beginning length was divided by the time it took for the envelope to completely disassemble.

#### Gaps in disassembling envelopes

Gaps appearing during the disassembly of the tau envelopes were counted manually. The number of gaps on each envelope were then divided by the envelope length at the start of the movie and by the length (in sec) of the movie.

#### Tau density on GDP-microtubules

The maximum intensity projection from the last 5 frames of the video was made to eliminate the impact of slight fluorescent intensity fluctuations on the intensity measurements. The mean intensity of tau signal on microtubules was measured from these maximum intensity projections, subtracted by the mean intensity of the background and divided by 5 (number of frames). Mean intensities were measured for all tested concentrations.

#### Kinesin landing rate

Kinesin-1 landings were counted manually from kymographs of the kinesin channel. Kinesin-1 landings were counted when landings visually appeared to be kinesin-1 molecules (based on fluorescence intensity and size).

#### Katanin-mediated microtubule disassembly rate

Katanin-mediated microtubule disassembly rate was measured as the change of envelope length over time. Tau envelopes were measured at their longest length and at the end of the captured movie (t=30 min). If a tau envelope disassembled completely before t=30 min, the disappearance of the fluorescence signal marked the end of the measurement. Envelope length was measured by manually drawing a line along the envelope in ImageJ and calculating the length in nm. Subsequently, the lengths were divided by time in sec and normalized to the length of the envelope prior to disassembly (at t=0 min).

#### Katanin severing in envelope

Katanin severing events were counted manually from fluorescence videos. Severing events were counted when a clear gap appeared in the tau-mCherry signal and disassembly of the microtubule was observed from the newly acquired boundaries.

#### Normalized tau density in envelopes after katanin addition

Tau density was measured at 5 timepoints after addition of katanin. First frame after katanin addition marks t=0 min. Tau density was measured within the envelope region, and normalized to the tau density within the same envelope at t=0 min.

### Live-cell experiments

#### Plasmids

Human tau sequence (h441-tau) N-terminally tagged with eGFP in pCDNA.4 vector was used as control tau. Tau sequence with deleted N terminus (tau 242-441) was created from control tau using one-step site-directed deletion^38^; primers Fwd: CGGCCGCACGCCTGCAGACAGCCCCCGTGCCCAT, Rev: GCAGGCGTGCGGCCGCGGCTCCGAATTCTTTGTATAGT). Human tubulin sequence (TUBA1B) fused with mScarlet was used for lentiviral and retroviral particles production. To increase the phosphorylation level, co-transfection was used with vectors pCDNA3 overexpressing Cdk5 and p25. For katanin overexpression, co-transfection of pLL vectors expressing katanin subunit p60 and GFP-tagged katanin subunit p80 was used.

#### Tau lysate preparation

HEK293T cells were co-transfected either with GFP-tau and empty pCDNA3 vector (1:1) or with GFP-tau and vectors pCDNA3 overexpressing Cdk5 and p25 (1:0.85:0.15) using linear Polyethylenimine (PEI; Polysciences, Inc.). Cells were harvested 48 hours after transfection by centrifugation and flash-frozen. Cell pellets were resuspended in 0.5 pellet volumes of lysis buffer (BRB80 supplemented with 1x phosphatase inhibitors (#4906845001, Sigma), 1x protease inhibitors (#04693159001, Sigma) and 0.05% Triton X-100 (# X100, Sigma)). The mixture was sonicated with three short pulses using the sonotrode MS1 (Hielscher Ultrasonics), setting “cycle” 1, “amplitude” 100% (30 kHz) on ice. The solution was transferred to 270 µl Beckman ultracentrifuge tubes and ultra-centrifuged in the Beckman 42.2 Ti rotor at 30000 x g, 4°C for 30 min in the Beckman Coulter Optima XPN-90 ultracentrifuge. The supernatant was directly used for experiments or flash-frozen in liquid nitrogen and stored at −80 °C. Lysate concentrations were measured at 488nm absorbance using NanoDrop ND-1000 spectrophotometer (Thermo Scientific).

#### Tau lysate imaging

Chambers were prepared as described before. Tau lysates were diluted 10x in AB buffer and added to surface-immobilized microtubules where tau envelope formation was captured for 3 min with 5 sec interval.

#### Elevated-pH treatment and live-cell imaging

IMCD-3 cells were transduced by lentiviral particles produced by co-transfection of HEK293T cells with lentiviral vector carrying the sequence for mScarlet-tubulin together with gag/pol and vsv-g vector (1:0.9:0.1). Medium containing lentiviral particles was collected 48 hours after transfection, filtered (0.45 µm pores), and used for transduction of IMCD-3 cells.

IMCD-3 cells expressing mScarlet tubulin were then transfected in OptiMEM media (Thermo Scientific) using Lipofectamine 2000 (Thermo Scientific) according to the manufacturer’s protocol. Cells were transfected in 8-well chambered coverslips (Ibidi), 0.5 µg DNA/well was used. Co-transfection of GFP-tau, Cdk5, and p25 was done in the ratio 1:0.85:0.15. The cells were grown in DMEM/F12 supplemented by FBS and Penicillin/Streptomycin (Thermo Scientific). The imaging was done 24 hours after transfection.

Cells were imaged every 20 sec for 10 min. After 1 min of imaging, the media (DMEM/F12 supplemented by FBS and Penicillin/Streptomycin) was changed for the regular media with pH adjusted to 8.4 with NaOH. Imaging was performed using TIRF microscope (Apo TIRF 60x Oil DIC N2; 488+561 exposure: 300 ms) using OKO-lab chamber (37°C, 5% CO_2_).

#### FRAP experiments

For FRAP experiments, the cells were prepared the same way as described in section ‘Elevated-pH treatment and live-cell imaging‘. A spinning-disk confocal microscope (Nikon CSU-W1) equiped with FRAP/photoactivation module was used to image and FRAP cells. Cells were imaged using CF Plan Apo VC 60XC WI objective (water immersion) and 488nm laser and FITC filter. Imaging was done on a single cell using 3 different settings: (1) the cell was imaged for 17 frames (100 ms exposure time, 500ms interval) before FRAP; (2) FRAP was performed on a circular region of 0.5 μm diameter; (3) the cell was imaged for 22 seconds directly after FRAP to visualize the recovery. The imaging was done at 37°C and 5% CO_2_.

#### Cell cycle experiments

For the monitoring of tau during cell cycle, U-2 OS human cell line (ATCC HTB-96) was transfected with the GFP-tau vector using the X-tremeGENE HP reagent (Sigma Aldrich) and then selected with 200 µg/mL zeocin. GFP-positive cells were sorted by fluorescent-activated cell sorting (BD FACS Aria Fusion). Cells were then transduced with retroviral particles containing the mScarlet-tubulin. Briefly, Platinum A cells were transfected with pMXs-Puro-mScarletI-Tubulinα, particles were collected after 48 hours and applied to cells, which were selected with 2.5 µg/mL puromycin. The resulting GFP-tau/mScarlet-tubulin cell line was grown on glass-bottom dishes in Fluorobrite medium with 10% FBS and glutamine and observed on a confocal Zeiss LSM 880 microscope at 37°C and 5% CO_2_.

#### Katanin experiment in cells – preparation

IMCD-3 cells were co-transfected with vectors for overexpression of mCherry-tau or mCherry, katanin subunit p60, katanin subunit p80-GFP, and Cdk5 and p25 or empty pCDNA3 vector (in the ratio: 1: 0.375: 0.375: 0.375: 0.375). 12 hours after transfection the cells were fixed using 4% PFA/PBS for 15 min followed by methanol at −20 °C for 2 min. Fixed cells were kept in PBS at 4 °C.

#### Katanin experiment in cells – immunostaining

Cells were blocked for 1 hour in 0.1% BSA/PBS and then stained with anti-β-tubulin antibody overnight (1:400; DSHB Hybridoma Product E7 deposited to the DSHB by Klymkowsky, M.^39^). After washing, cells were incubated with anti-mouse secondary antibody conjugated with Alexa-647 (Thermo Scientific). Cells were captured using a confocal microscope Leica Stellaris 8 (HC PL APO CS2 63x/1.40 OIL, WLL laser).

### Live-cell image analysis

*Tau lysate analysis*. Microtubule lengths were measured by using the IRM signal, tau envelope lengths were measured by using the fluorescent tau signal after 3 min of incubation. The envelope coverage represents the sum of all tau envelopes lengths divided by the sum of all microtubule lengths within one field of view.

#### Coefficient of variation (CoV)

To determine the CoV in the elevated-pH treatment experiments, a circle was drawn inside the cell, covering (most of) the area of the cell (in case of large cells, two circular areas were averaged). For monitoring GFP-tau during the cell cycle, whole cells were manually selected. The CoV was determined using ImageJ from the standard deviation of the tau fluorescent signal within the ROI, divided by the mean. In elevated-pH treatment experiment, coV was analyzed before, at t = 0 min and t = 9 min after elevated-pH treatment.

#### Microtubule CoV

The microtubule CoV was determined as the regular CoV, however, the ROI in this case was a line drawn on a single microtubule inside the cell. For every cell, 3 random microtubules were measured and the CoV was averaged.

#### Pearson’s R

Cells in interphase or in mitosis were manually selected based on the presence or absence of a mitotic spindle, and Pearson’s correlation coefficient between the GFP-tau and the mScarlet-tubulin channels was calculated with the Coloc 2 plug-in in ImageJ.

#### Tau density on microtubules

To show that the affinity of tau to the microtubule is different in different cell groups, the mean intensity of tau on the microtubule was measured before elevated-pH treatment and divided by the mean intensity of the same-size region in the cytoplasm next to the measured microtubule at the same timepoint. 25 cells were analyzed (average of 5 microtubules was used for each cell).

#### Mean intensity of tau in the cell

To show that the transfection of tau was comparable between the groups, the mean intensity of the GFP signal was measured in random circular regions of the cytoplasm before elevated-pH treatment.

#### Normalized tau density in patches

To show that the density of tau in patches is stable over time for the tau and tau-Cdk5 group, while the density of tau-ΔN is increasing, the intensity of tau in the patches at 5 different timepoints after elevated-pH treatment was analyzed. For tau-ΔN the intensity over the entire length of the microtubule was analyzed. The mean intensity in a tau positive region on the microtubule (tau patch or full microtubule) was measured and subtracted by the mean intensity of the same-size region next to the microtubule. All timepoints were normalized to the intensity of tau before the elevated-pH treatment. 10 cells were analyzed and at each timepoint the average tau density on 5 microtubules was used.

#### Normalized tau density in the cell

To quantify the recovery of tau signal, the mean density of tau signal on the microtubules was analyzed during the time course. The mean tau intensity in the cell (ROI comprising most of the cell) was measured at all time points and subtracted by the intensity of the GFP signal in the cytoplasm next to the microtubules). The intensity in each time point was then normalized to the intensity before elevated-pH treatment. Both curves (control tau cells and tau-Cdk5 cells) were fitted using Matlab fitting tool using exponential recovery curve: f(x)= a-b*exp(-c*x) with exponential time constant 1/c [min].

#### FRAP analysis – recovery time constant

To analyze the recovery of the GFP signal on the microtubule after photobleaching, the intensity of the GFP signal in a small ROI on the microtubule in the bleached region of the cell was analyzed using ImageJ. The same-size region on the microtubule in the same cell but outside the bleached area was used as a reference (ref) and the same-size region outside the cell was used as a background (bg). 14-15 cells were analyzed. The curves were double-normalized according to the equation: F*frap-normalized*(*t*) = (F*ref-pre*/[F*ref*(*t*) - F*bg*(*t*)])([F*frap*(*t*) - F*bg*(*t*)]/F*frap-pre*), where F*ref-pre = Σ(t=0;t=17) ([Fref(t) - Fbg(t)]/fprebleach);* F*frap-pre = Σ(t=0;t=17) ([Ffrap(t) - Fbg(t)]/fprebleach); fprebleach = 17;* F*ref*(*t*) is the reference fluorescence intensity on the microtubule in the same cell but not in the bleached region; F*_frap_*(*t*) is the fluorescence intensity on the microtubule in the bleached ROI; F*_bg_*(*t*) is the fluorescence intensity in a background ROI outside the cells; F*_ref-pre_*is the mean fluorescence intensity of the reference ROI before the bleaching after background subtraction; F*_frap-pre_* is the mean fluorescence intensity of the bleached ROI before the bleaching after background subtraction. The normalized data was fitted using the Matlab fitting tool using the equation: *y = a*exp(-b*x)+c;* where *b* = rate constant and *c* = asymptote.

#### FRAP analysis – immobile fraction

To calculate the percentage of immobile fraction, we used the equation: Immob = [1-(*c*-*F*_0_)/(1-*F*_0_)]*100; where *c* is the asymptote (of the fitted curve), and *F_0_* is the normalized intensity immediately after the bleaching.

#### Katanin experiment – relative tubulin and katanin density

The mean density of stained tubulin signal in the transfected cell relative to the mean density of surrounding cells non-transfected with katanin was analyzed using ImageJ (average tubulin intensity of 3 circular ROIs in the cytoplasm of the transfected cell divided by the average tubulin intensity in 3 circular ROIs in 3 randomly selected cells in close vicinity to the analyzed cell). The relative tubulin density was either correlated to the relative density of the katanin signal (correlation plot with the linear regression) or plotted according to the experimental groups in a scatter plot (for this purpose only cells with relative intensity of katanin between 0.5-3 were plotted).

### Mass Spectrometry

Samples were analyzed using a liquid chromatography system Agilent 1200 (Agilent Technologies) connected to the timsToF Pro PASEF mass spectrometer equipped with Captive spray (Bruker Daltonics). Mass spectrometer was operated in a positive data-dependent mode. Five microliters of peptide mixture were injected by autosampler on the C18 trap column (UHPLC Fully Porous Polar C18 2.1mm ID, Phenomenex). After 5 min of trapping at a flow rate of 20 µL/min, peptides were eluted from the trap column and separated on a C18 column (Luna Omega 3 μm Polar C18 100 Å, 150 x 0.3 mm, Phenomenex) by a linear 35 min water−acetonitrile gradient from 5% (v/v) to 35% (v/v) acetonitrile at a flow rate of 4 µL/min. The trap and analytical columns were both heated to 50°C. Parameters from the standard proteomics PASEF method were used to set timsTOF Pro. The target intensity per individual PASEF precursor was set to 6000, and the intensity threshold was set to 1500. The scan range was set between 0.6 and 1.6 V s/ cm^2^ with a ramp time of 100 ms. The number of PASEF MS/MS scans was 10. Precursor ions in the m/z range between 100 and 1700 with charge states ≥2+ and ≤6+ were selected for fragmentation. The active exclusion was enabled for 0.4 min. The raw data were processed by PeaksStudio 10.0 software (Bioinformatics Solutions, Canada). The search parameters were set as follows: enzyme – trypsin (specific), carbamidomethylation as a fixed modification, oxidation of methionine, phosphorylation (STY) and acetylation of protein N-terminus as variable modifications.

### Mass spectrometry sample preparation

#### Tau samples

Tau with different levels of phosphorylation (phospho-tau, dephospho-tau, Bact-tau, Bact-tau-Cdk5) was prepared as stated above.

#### Spin down samples

1μM phospho-tau (insect cell expressed) was diluted in Mass Spec buffer (50mM HEPES, 75mM KCl, 10 µM taxol, 10 mM dithiothreitol, 1 mM Mg-ATP) and added to taxol-stabilized microtubules to reach a final volume of 100 µl. Microtubules and tau were incubated for 10 min at room temp and centrifuged for 30 min at 18000 x g in a Microfuge 18 Centrifuge (Beckman Coulter). After centrifugation the supernatant was separated from the pellet and used as the sample indicated by the ‘tau in solution‘. The pellet was resuspended in a four-fold lower volume of Mass Spec buffer and centrifuged again to ensure a more homogeneous sample. After the second centrifugation, the supernatant was discarded and the pellet was resuspended in BRB80 and used as the sample indicated by the ‘tau in envelopes‘.

### Mass Spectrometry analysis

#### Phosphorylation degree

For each phosphorylation site, all peptides were studied that include the specific site, phosphorylated or not. Each peptide is found with a relative intensity, that gives an indication of the density at which that peptide is detected in the sample. Each peptide can be found with the specific phosphorylation site phosphorylated or not. The sum of the relative intensities of the peptide in phosphorylated state was divided by the sum of the relative intensities of the peptide in non-phosphorylated state to obtain the phosphorylation degree of the phosphorylation site. Each tau sample was prepared as triplicates and each individual sample was analyzed separately, resulting in 3 individual phosphorylation degree values for each phosphorylation site. The graphs in the paper display the mean ± s.d. for each sample at each phosphorylation site, either plotted along the amino acid sequence of tau, or as individual phosphorylation sites.

#### Total relative intensity

For each phosphorylation site, all relative intensities at which the peptides covering the specific phosphorylation were summed up to get the total relative intensity.

### Statistics and reproducibility

For representative plots and figures, whenever not specifically stated in the caption, all data was collected from at least three independent trials. All repeated independent experiments showed similar results and no data was excluded from the manuscript. Unless stated otherwise, all data were analyzed manually using ImageJ (FIJI) or Matlab (R2020b). Graphs were created using Matlab R2020b and statistical analyses were performed using the same software. Major points on graphs represent data means and the error bars represent variation or associated estimates of uncertainty.

### Data availability

Source data files for all figures and supplementary figures, are available with this manuscript.

## Acknowledgements

We thank the members of the Lansky-Braun lab and Balastik lab for their feedback and helpful discussion and T. Šmídová and K. Konečná for their technical support. This work was supported by Czech Science Foundation grant 19–27477X to Z.L. and L.L., 21-24571S to R.W. and 23-07703S to M.Br., the project National Institute for Neurological Research (Programme EXCELES, ID LX22NPO5107) - Funded by the European Union - Next Generation EU, the Charles University Grant Agency (GAUK no. 373821 to V.S.), the project ‘Grant Schemes at CU’ (reg. no. CZ.02.2.69/0.0/0.0/19_073/0016935) to V.S. and V.D and by Czech Health Research Council grant no. NV18-04-00085 to M.Ba. We acknowledge the Imaging Methods Core Facility at BIOCEV, institution supported by the MEYS CR (LM2023050 Czech-BioImaging) for their support & assistance in this work, the CF Protein Production of CIISB, Instruct-CZ Centre, supported by MEYS CR (LM2023042) for protein production, CF Structural mass spectometry of CIISB, Instruct-CZ Centre, supported by MEYS CR (LM2018127) and European Regional Development Fund-Project „UP CIISB“ (No. CZ.02.1.01/0.0/0.0/18_046/0015974) for mass spectometry sample analysis, the microscope facility of FGU, supported by MEYS CR (Large RI Project LM2018129 Czech-BioImaging) and ERDF (project No. CZ.02.1.01/0.0/0.0/18_046/0016045) and Vinicna Microscopy Core Facility co-financed by the Czech-BioImaging large RI project LM2023050 for their support and assistance in microscopy. Computational resources were supplied by the project “e-Infrastruktura CZ” (e-INFRA LM2018140) provided within the program Projects of Large Research, Development and Innovations Infrastructures.

## Author contributions

The manuscript was conceptualized by M.Br., M.Ba., Z.L.; methods were developed by V.S., R.W., E. L., A.K., V.D.; TIRF in vitro experiments were performed by V.S., E. L., A.K.; tau lysate preparation and experiments were performed by R.W., E.L.; live-cell experiments were performed by R.W., V.D.; FRAP experiments were performed and optimized by V.S., R.W.; data were analyzed by V.S., R.W., E.L., A.K., V.D.; resources were provided by V.H., C.J.; the manuscript was written by V.S., M.Br., Z.L., with reviewing and editing by R.W., M.Ba.; visualization was done by V.S.; the project was supervised by L.L., M.Br., M.Ba., Z.L.; funding was acquired by V.S., R.W., V.D., L.L., M.Br., M.Ba., and Z.L.

## Competing interests

The authors declare no competing interests.

## Supplemental information

### Supplementary movie captions

**Movie 1** related to Fig. 1c: **1.5nM phospho-tau on taxol-stabilized microtubules.**

A time-lapse movie of 1.5 nM phospho-tau-GFP (magenta) added to surface-immobilized taxol-stabilized microtubules and imaged for 5 minutes.

**Movie 2** related to Fig. 1c: **1.5nM dephospho-tau on taxol-stabilized microtubules.**

A time-lapse movie of 1.5 nM dephospho-tau-GFP (cyan) added to surface-immobilized taxol-stabilized microtubules and imaged for 5 minutes.

**Movie 3** related to Fig. 1e: **10nM Bact-tau in presence of active Cdk5.**

A time-lapse movie of 10 nM Bact-tau-GFP (magenta) on surface-immobilized taxol-stabilized microtubules (black). Active Cdk5/p35 was added to the measurement chamber at t=0 min.

**Movie 4** related to Fig. 1e: **10nM Bact-tau in presence of deactivated Cdk5.**

A time-lapse movie of 10 nM Bact-tau-GFP (cyan) on surface-immobilized taxol-stabilized microtubules (black). Deactivated Cdk5/p35 (control) was added to the measurement chamber at t=0 min.

**Movie 5** related to Fig. 2g: **10nM Bact-tau removal in presence of active Cdk5.**

A time-lapse movie of 10 nM Bact-tau-GFP (magenta) on surface-immobilized taxol-stabilized microtubules (black). Bact-tau was removed from solution in presence of active Cdk5/p35.

**Movie 6** related to Fig. 2g: **10nM Bact-tau removal in presence of deactivated Cdk5.**

A time-lapse movie of 10 nM Bact-tau-GFP (cyan) on surface-immobilized taxol-stabilized microtubules (black). Bact-tau was removed from solution in presence of deactivated Cdk5/p35.

**Movie 7** related to Fig. 3a: **FRAP of tau-GFP (control) cell.**

A time-lapse movie showing an IMCD-3 cell overexpressing GFP-tau (control, cyan) on which FRAP is performed at t=0 min. GFP-tau signal was monitored for 10 seconds before FRAP and 20 seconds after FRAP.

**Movie 8** related to Fig. 3a: **FRAP of tau-GFP-deltaN cell.**

A time-lapse movie showing an IMCD-3 cell overexpressing GFP-tau-ΔN (tau-ΔN) on which FRAP is performed at t=0 min. GFP-tau signal was monitored for 10 seconds before FRAP and 20 seconds after FRAP.

**Movie 9** related to Fig. 3a: **FRAP of tau-GFP-Cdk5 cell.**

A time-lapse movie showing an IMCD-3 cell overexpressing GFP-tau-Cdk5/p25 (tau-Cdk5) on which FRAP is performed at t=0 min. GFP-tau signal was monitored for 10 seconds before FRAP and 20 seconds after FRAP.

**Movie 10** related to Supplementary Fig. 3g (left): **pH treatment on tau (control) cell.**

A time-lapse movie showing an IMCD-3 cell overexpressing GFP-tau (control) on which elevated-pH treatment is performed at t=0 min. GFP-tau signal (left) and mScarlet-tubulin signal (right) were monitored for 10 minutes; 1 min before treatment and 9 minutes after treatment.

**Movie 11** related to Supplementary Fig. 3g (middle): **pH treatment on tau-ΔN cell.**

A time-lapse movie showing an IMCD-3 cell overexpressing GFP-tau-ΔN (tau-ΔN) on which elevated-pH treatment is performed at t=0 min. GFP-tau signal (left) and mScarlet-tubulin signal (right) were monitored for 10 minutes; 1 min before treatment and 9 minutes after treatment.

**Movie 12** related to Supplementary Fig. 3g (right): **pH treatment on tau-Cdk5 cell.**

A time-lapse movie showing an IMCD-3 cell overexpressing GFP-tau-Cdk5/p25 (tau-Cdk5) on which elevated-pH treatment is performed at t=0 min. GFP-tau signal (left) and mScarlet-tubulin signal (right) were monitored for 10 minutes; 1 min before treatment and 9 minutes after treatment.

**Movie 13** related to Supplementary Fig. 4a (left): **Kinesin-1 molecules walking on microtubules in presence of phospho-tau envelopes.**

A time-lapse movie showing kinesin-1-GFP molecules (white) walking on taxol-stabilized microtubules partly covered by phospho-tau envelopes (magenta).

**Movie 14** related to Supplementary Fig. 4a (right): **Kinesin-1 molecules walking on microtubules in presence of dephospho-tau envelopes.**

A time-lapse movie showing kinesin-1-GFP molecules (white) walking on taxol-stabilized microtubules partly covered by dephospho-tau envelopes (cyan).

**Movie 15** related to Fig. 4a: **Katanin severing microtubules covered with phospho-tau envelopes.**

A time-lapse movie showing katanin-GFP (yellow) severing taxol-stabilized microtubules partly covered by phospho-tau envelopes (magenta).

### Supplementary figures

**Supplementary Fig. 1:**
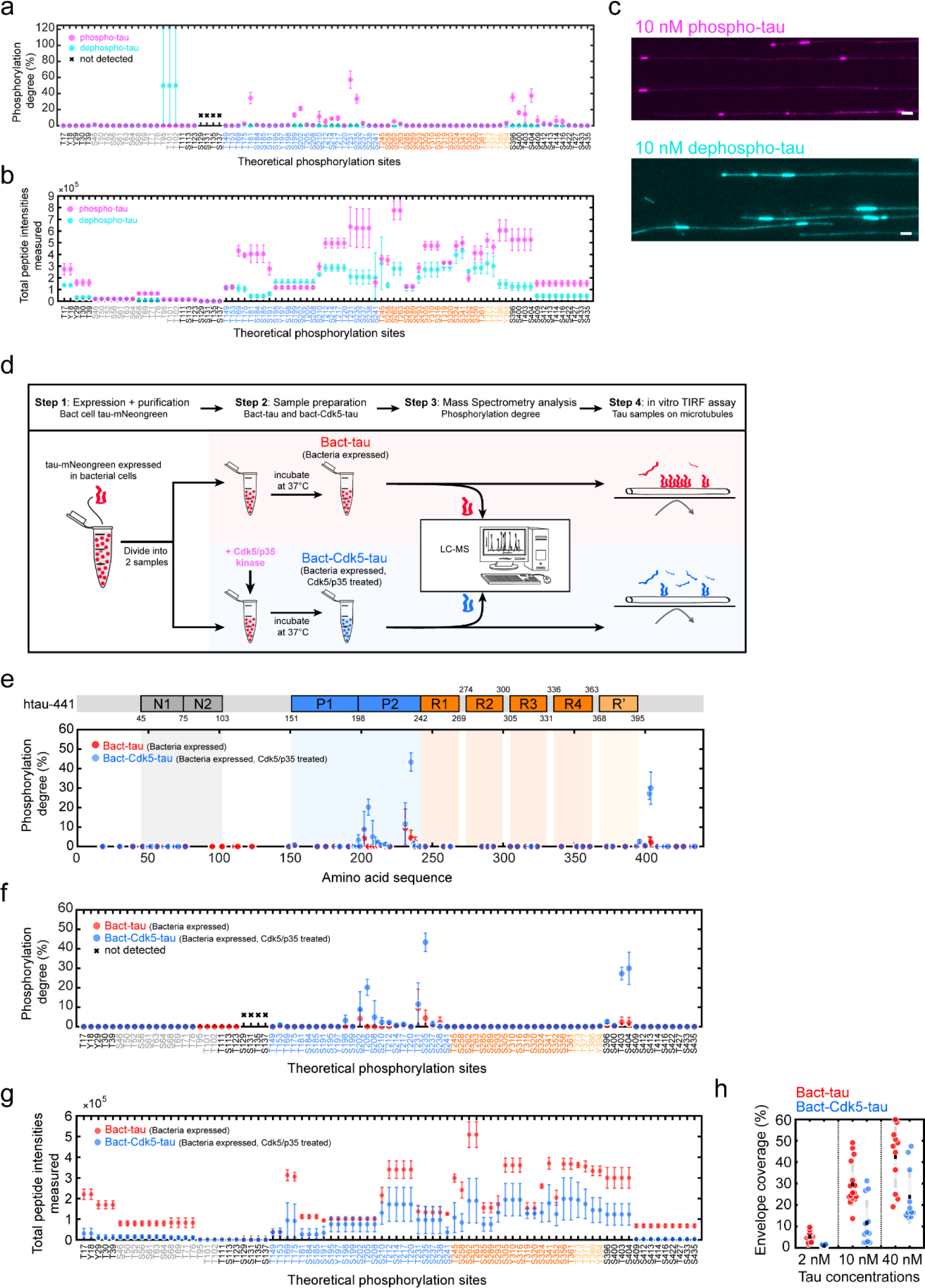
**a**. Mass-spectrometry-determined degree of phosphorylation of phospho-tau (magenta) and dephospho-tau (cyan). Phosphorylation degrees are presented as mean ± s.d. for each tau sample. If no peptides were detected that covered a specific phosphorylation site, these phosphorylation sites are marked with a black cross. The color-coded legend corresponds to the domains along the tau sequence highlighted in Fig. 1b. **b**. Total relative intensities measured for all peptides covering the specific phosphorylation site, corresponding to data in a and Fig. 1b. Relative intensities are presented as mean ± s.d. for each tau sample. The color-coded legend corresponds to the domains along the tau sequence as shown in Fig. 1b. **c**. Fluorescence micrographs of 10 nM phospho-tau (left, phospho-tau in magenta) or 10 nM dephospho-tau (right, dephospho-tau in cyan) on surface-immobilized microtubules after 3 min of incubation. Scale bars: 2 μm. **d**. Schematics of sample preparation of bacterial expressed tau-mNeongreen (red), and Cdk5/p35-treated bacterial expressed tau-mNeongreen (blue). **e**. Mass-spectrometry-determined degree of phosphorylation of Bact-tau (red) and Bact-Cdk5-tau (blue). Phosphorylation degree is presented as mean ± s.d. and displayed at the location of the phosphorylation site along the amino acid sequence (schematic of the sequence is shown above the plot). The domains on the tau sequence are color-coded: N-terminal domains (N1, N2, grey), proline-rich domains (P1, P2, blue), microtubule-binding repeats (R1-R4, orange), and the domain pseudo-repeat (R‘, light orange). **f**. Mass-spectrometry-determined degree of phosphorylation of Bact-tau or Bact-Cdk5-tau for individual phosphorylation sites. Phosphorylation degree is presented as mean ± s.d. If no peptides were detected that covered a specific phosphorylation site, these phosphorylation sites are marked with a black cross. The color-coded legend corresponds to the domains along the tau sequence in Fig. 1e. **g**. Total relative intensities measured for all peptides covering the specific phosphorylation site, corresponding to data in e and f. Relative intensities are presented as mean ± s.d. The color-coded legend corresponds to the domains along the tau sequence as shown in Fig. 1e. **h**. Percentage of taxol-stabilized microtubules covered with a tau envelopes after 3 min incubation. Envelope coverage for Bact-tau was 5.1 ± 2.5% at 2 nM, 29.6 ± 10.1% at 10 nM, and 42.4 ± 14.4% at 40 nM (mean ± s.d., each data point represents a single field of view, n=12, 16, 11 independent experiments). Envelope coverage for Bact-Cdk5-tau was 0.9 ± 0.5% at 2nM, 11.7 ± 10.7% at 10nM, and 23.8 ± 12.6% at 40nM (mean ± s.d., each data point represents a single field of view, n=12, 12, 11 independent experiments).

**Supplementary Fig. 2:**
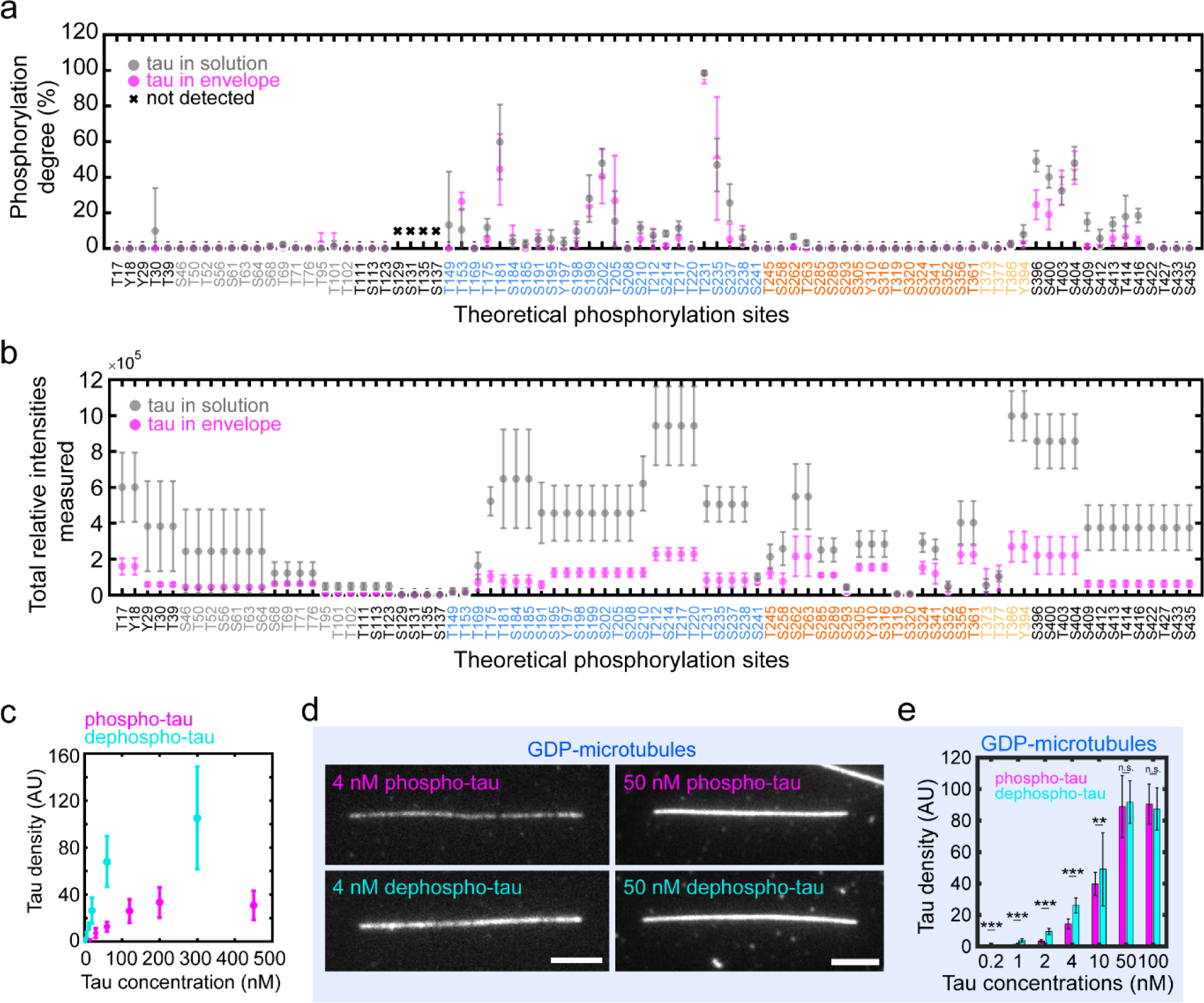
**a**. Mass-spectrometry-determined phosphorylation degree of spin down sample, where unbound tau was found in the supernatant (tau in solution, grey), and cooperatively bound tau found in the pellet (tau in envelope, magenta). Phosphorylation degrees are displayed as mean ± s.d. If no peptides were detected that covered a specific phosphorylation site, these phosphorylation sites are marked with a black cross. The color-coded legend corresponds to the domains along the tau sequence as shown in Fig. 2b. **b.** Total relative intensities of all peptides that covered the specific phosphorylation site, corresponding to data in a and Fig. 2b. Intensities are displayed as mean ± s.d. for unbound tau from the supernatant (tau in solution, grey), and cooperatively bound tau from the pellet (tau in envelope, magenta). The color-coded legend corresponds to the domains along the tau sequence as shown in Fig. 2b. **c**. Density of phospho-tau (magenta) or dephosho-tau (cyan) measured on GMPCPP microtubules as a function of the concentration of phospho-tau or dephosho-tau in solution (phospho-tau: n=12 fields of view in 4 independent experiments; dephospho-tau: n=9 fields of view in 3 independent experiments). **d.** Fluorescence micrographs of phospho-tau (magenta) or dephospho-tau (cyan) on glycerol-stabilized GDP-microtubules after 1 min of incubation. Scale bars: 2 μm. **e.** Concentration (nM) of phospho-tau (magenta) and dephospho-tau (cyan) plotted against the tau density (AU) on GDP-microtubules. For 0.2 nM: 0.07 ± 0.21 (phospho-tau) and 0.37 ± 0.20 (dephospho-tau) (mean ± s.d., n=36 microtubules in N=2 independent experiments, two-sided t-test: p=1.31*10^-8^); 1 nM: 0.91 ± 0.36 (phospho-tau) and 3.75 ± 1.09 (dephospho-tau) (mean ± s.d., n=36 microtubules in N=2 independent experiments, two-sided t-test: p=7.81*10^-26^); 2 nM: 3.43 ± 0.89 (phospho-tau) and 9.51 ± 1.92 (dephospho-tau) (mean ± s.d., n=36 microtubules in N=2 independent experiments, two-sided t-test: p=2.53*10^-33^); 4 nM: 14.35 ± 3.10 (phospho-tau) and 26.12 ± 4.79 (dephospho-tau) (mean ± s.d., n=36 microtubules in N=2 independent experiments, two-sided t-test: p=3.29*10^-28^); 10 nM: 39.71 ± 7.38 (phospho-tau) and 49.12 ± 23.25 (dephospho-tau) (mean ± s.d., n=51 and 57 microtubules respectively in N=3 independent experiments, two-sided t-test: p=0.0048); 50 nM: 88.88 ± 19.70 (phospho-tau) and 91.69 ± 13.45 (dephospho-tau) (mean ± s.d., n=45 and 44 microtubules respectively in N=3 independent experiments, two-sided t-test: p=0.3939); 100 nM: 90.43 ± 12.77 (phospho-tau) and 87.40 ± 13.38 (dephospho-tau) (mean ± s.d., n=20 and 30 microtubules respectively in N=1 independent experiment, two-sided t-test: p=0.4397).

**Supplementary Fig. 3:**
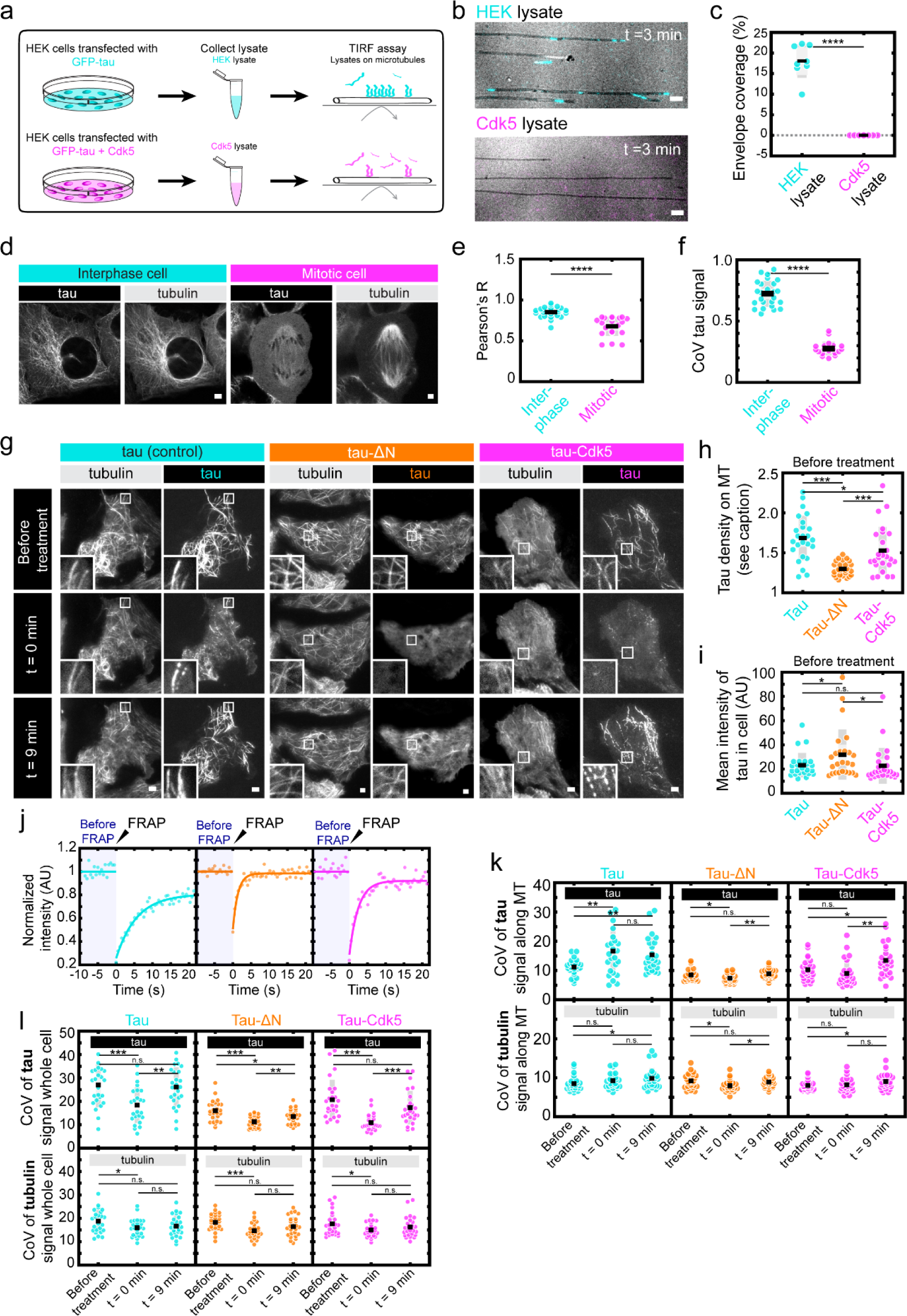
**a**. Schematics of preparation of lysates from HEK cells transfected with GFP-tau (HEK lysate, top, cyan) and of lysate prepared from HEK cells transfected with GFP-tau and Cdk5/p25 (Cdk5 lysate, bottom, magenta). **b**. Multichannel fluorescence micrographs of HEK lysate (top, cyan) and Cdk5 lysate (bottom, magenta) added to surface immobilized taxol-stabilized microtubules (black) after 3 min of incubation. Scale bars: 2 μm. **c**. Percentage of taxol-stabilized microtubules covered by a tau envelope (envelope coverage) after addition of HEK lysate was 18.2 ± 3.4% (mean ± s.d., n = 8 fields of view in 8 independent experiments), and after addition of Cdk5 lysate was 0.0 ± 0.0% (mean ± s.d., n = 8 fields of view in 4 independent experiments). Two-sided t-test, p=4.3821*10^-10^. **d**. Fluorescence micrographs of GFP-tau (left) and mScarlet-tubulin (right) in U-2 OS cells at different phases of the cell cycle. Cells in interphase (left panels, cyan) show higher binding of tau to microtubules compared to cells in mitosis (right panels, magenta). **e**. Pearson’s R correlation coefficient between the GFP-tau and mScarlet-tubulin signal differs significantly between interphase and mitotic cells. Pearson’s R in interphase cells was 0.85 ± 0.06 and in mitotic cells 0.68 ± 0.12 (mean ± s.d., interphase, n=25 cells in 3 independent experiments; mitosis, n=20 cells in 3 independent experiments). Two-sided t-test, p=1.4224*10^-7^. **f**. Coefficient of variation (CoV) of the tau signal measured over the whole cell reflecting the difference between tau signal on microtubules and in cytoplasm. CoV of tau signal in interphase cells was 0.72 ± 0.11 and in mitotic cells 0.28 ± 0.06 (mean ± s.d., interphase n=25 cells in 3 independent experiments, mitosis n=19 cells in 3 independent experiments). Two-sided t-test, p=5.2528e-20. **g.** Fluorescence micrographs of cells before elevated-pH treatment (left), at t=0 min after elevated-pH treatment (middle) and t=9 min after elevated-pH treatment (right). 3 types of cells were subjected to elevated-pH treatment; control GFP-tau cells (tau, cyan), GFP-tau-ΔN cells (tau-ΔN, orange), and GFP-tau-Cdk5 cells (tau-Cdk5, magenta). Top panels represent the GFP-tau channel, bottom panels the mScarlet-tubulin channel. Scale bars: 10μm. **h.** Density of tau on microtubules before elevated-pH treatment, compared to density of tau in the cytoplasm, to give a measure of the binding affinity of tau in the different cells. Tau density on microtubules compared to cytoplasm in tau cells was 1.68 ± 0.28, in tau-ΔN cells was 1.30 ± 0.09, in tau-Cdk5 cells was 1.53 ± 0.30 (mean ± s.d., n=25 cells for each group in 4 independent experiments). Two-sided t-test p-values (left-to-right): 3.26*10^-8^, 0.0594, 5.89*10^-4^. **i.** Mean intensity of tau over the whole cell, as an indication of the expression level. Mean intensity over the wholle cell in tau cells was 23.2 ± 10.1, in tau-ΔN cells was 31.9 ± 20.8, and in tau-Cdk5 cells was 22.8 ± 14.7 (mean ± s.d., n=25 cells for each group in 4 independent experiments). Two-sided t-test p-values (left-to-right): 0.0658, 0.8901, 0.0774. **j**. Fluorescence recovery curves after FRAP for control GFP-tau cells (left, cyan); GFP-tau-ΔN cells (middle, orange); GFP-tau-Cdk5 cells (right, magenta). Normalized tau intensity within the FRAP area is plotted over time. The shaded blue area marks the period before FRAP, the black arrow indicates the timepoint at which FRAP occurred. **k.** Coefficient of variation of GFP-tau signal (top) or mScarlet-tubulin signal (bottom) measured along the microtubule lattice in GFP-tau (cyan), GFP-tau-ΔN (orange), and GFP-tau-Cdk5 cells (magenta). Coefficient of variation was measured at 3 timepoints: before elevated-pH treatment (left), at t=0 min after elevated-pH treatment (middle), and at t=9 min after elevated-pH treatment (right). **l.** Coefficient of variation of GFP-tau signal (top) or mScarlet-tubulin signal (bottom) measured over the whole cell in GFP-tau (cyan), GFP-tau-ΔN (orange), and GFP-tau-Cdk5 cells (magenta). Coefficient of variation was measured at three timepoints: before elevated-pH treatment (left), at t=0 min after elevated-pH treatment (middle), and t=9 min after elevated-pH treatment (right).

**Supplementary Fig. 4:**
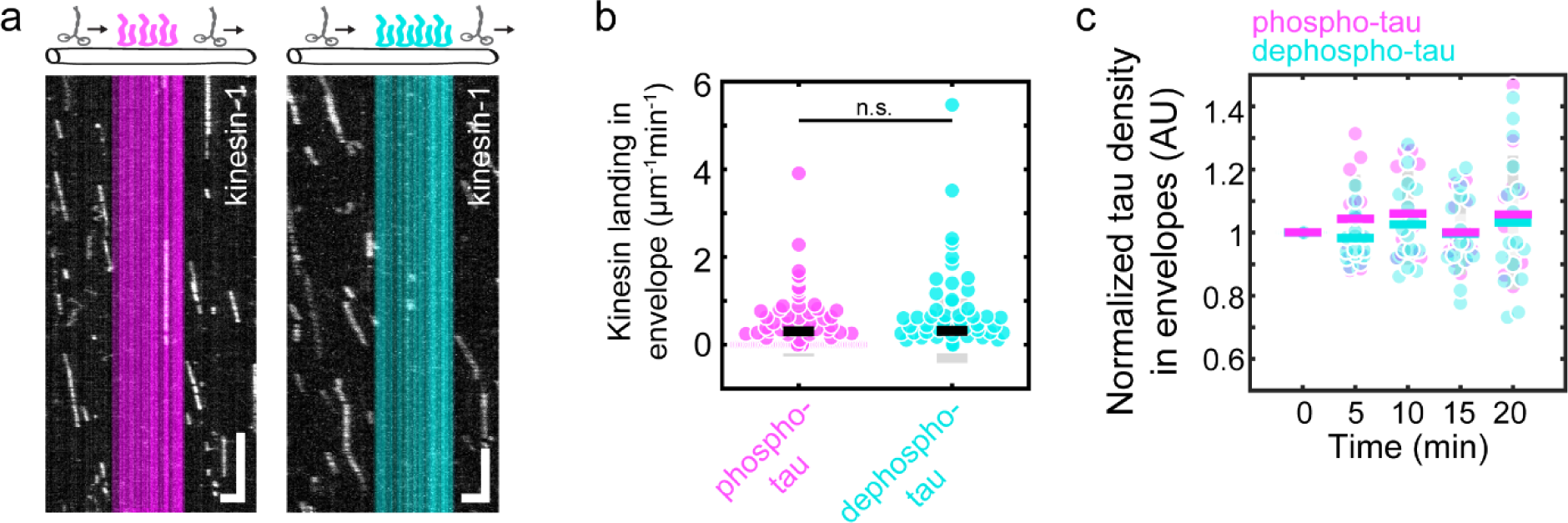
**a**. Fluorescence kymographs showing kinesin-1-GFP molecules (white) moving processively outside tau envelope region, independent of the phosphorylation degree of tau molecules forming the envelope (phospho-tau, left, magenta; dephospho-tau, right, cyan). Kinesin-1 landing but no processive movement was observed within the envelope boundaries. Scale bars: 2 μm, 1 s. **b**. Kinesin-1 landing rate within phospho-tau envelopes was 0.31 ± 0.58 µm^-1^min^-1^ and within dephospho-tau envelopes was 0.32 ± 0.73 µm^-1^min^-1^ (mean ± s.d., phospho-tau: n=100 envelopes in 5 experiments; dephospho-tau: n=125 envelopes in 5 experiments). Two-sided t-test, p=0.8894. **c**. Normalized tau density in envelopes after katanin addition for dephospho-tau was 0.98 ± 0.07, 1.02 ± 0.13, 1.00 ± 0.13, 1.03 ± 0.21 for 5, 10, 15, 20 minutes, respectively (mean ± s.d., n=16 envelopes in 2 individual experiments). Normalized tau density for phospho-tau was 1.04 ± 0.14, 1.06 ± 0.13, 1.00 ± 0.10, 1.06 ± 0.18 for 5, 10, 15, 20 minutes, respectively (mean ± s.d., n=13 envelopes in 4 individual experiments).

## Notes

### Competing Interest Statement

The authors have declared no competing interest.

